# Signal Transformations and New Timing Rules of Hippocampal CA3 to CA1 Synapses

**DOI:** 10.1101/2022.05.26.493588

**Authors:** Sandra Gattas, Aliza A. Le, Javad Karimi Abadchi, Ben Pruess, Yanning Shen, A. Swindlehurst, Michael A. Yassa, Gary S. Lynch

## Abstract

The synapse is the fundamental unit of communication in the nervous system. Determining how information is transferred across the synaptic interface is one of the most complex endeavors in neuroscience, owing to the large number of contributing factors and events. An approach to solving this problem involves collapsing across these complexities to derive concise mathematical formulas that fully capture the governing dynamics of synaptic transmission. We investigated the feasibility of deriving such a formula – an input-output transformation function for the CA3 to CA1 node of the hippocampus -- using the Volterra expansion technique for nonlinear system identification. The entirety of the field EPSP in the apical dendrites of mouse brain slices was described with >94% accuracy by a 2^nd^ order equation that captured the linear and nonlinear influence of past inputs on current outputs. This function generalized to cases not included in its derivation and uncovered previously undetected timing rules. The basal dendrites expressed a substantially different transfer function and evidence was obtained that, unlike the apical system, a 3^rd^ order system or higher will be needed for complete characterization. Collectively, these results describe a readily implemented and unusually sensitive means for evaluating the effects of pharmacological treatments and disease related conditions on synaptic dynamics. At scale, the approach will also provide information needed for the construction of biologically realistic models of brain networks.

## Introduction

Mathematical characterization of the composite synaptic response generated by small collections of input axons is a critical step in understanding communication among brain subsystems. Efforts in this direction encounter a series of challenges. The cortical telencephalon operates with a diverse set of signaling patterns embodied in spike patterns, which requires synaptic descriptions to include a very large number of input possibilities. Relatedly, such temporal patterns engage an extraordinarily diverse array of cellular and concomitant physiological events with different time courses on both sides of the synapse [1–8]. Past responses generated in response to past inputs thus have a strong influence on current ones as exemplified in the well-studied frequency facilitation effect [9, 10]. The description problem is further complicated by the presence of feedforward inhibitory neurons that are engaged by an active input and innervate the target dendrites for that input. This results in responses that are admixtures of monosynaptic and disynaptic events with the latter acting to shunt currents generated by the former [11]. In all, the seemingly simple potentials recorded in conventional experiments using repetitive stimulation reflect a large array of interacting variables.

The present study investigated the feasibility of capturing the size and waveform of synaptic responses in a concise mathematical formulation that accommodates time-varying input patterns. The experimental strategy followed an engineering approach referred to as ‘system identification’ that aims at accurately characterizing all possible operations performed by an unknown system (i.e., a black box). Work of this kind, using inputs sufficient to tap into the system’s full operational range, shows that it is possible to obtain a unique solution for input-output transformations – a single formula or transfer function that predicts outputs to arbitrary inputs – for electronic circuits. System identification has also been used to capture dynamics of different biological systems and across species [12]. The experimental question was thus whether system identification can also be used to describe axo-dendritic synaptic dynamics despite the immense complexity of such a biological system.

Beyond its importance for characterizing the manner in which variations in afferent patterns are differentially translated into dendritic output, an accurate transfer function would provide a novel and precise means for addressing widely studied issues in neuroscience. By design, the approach samples the entirety of responses rather than single measures such as slope or amplitude. This along with the above noted use of a full range of time-varying inputs, as opposed to stereotyped stimulation patterns, result in a new level of sensitivity for assessing the effects of pharmacological and genetic manipulations. A predictive formula would as well enable quantitative tests for differences between synaptic populations in the processing of realistic input patterns, an issue that is of vital importance to the construction of network models. The work reported here accordingly attempted to develop transfer functions for both the apical and basal dendrites of hippocampal field CA1, one of the most intensively studied brain regions.

## Results

### Signal transformation in CA1 apical dendrites

#### Using Volterra series expansion technique to identify the CA1 apical dendritic operations

Characterizing a system’s operations (i.e., identifying its transfer function) differs depending on whether the system is linear or nonlinear. For a linear, time-invariant system, estimating the impulse response function *h(i)* would fully characterize the system, but a nonlinear system requires higher order terms (h(k,m),.., h(k,m,..,p), **Fig. 1A**). We identified, up to the second order, the CA1 apical dendritic system transfer function by estimating both the first and second order kernels *h(i)* and *h(k,m)*, respectively, to capture linear and nonlinear dynamics, respectively, using the Volterra Series Expansion (VSE) for system identification [12]. This general form of the VSE fully captures n^th^ order system dynamics in an agnostic manner, with no prior assumptions about the system, and thereby avoids the generation of partially complete or misleading models [12]. Here, we have truncated the series expansion by making two assumptions: 1) system order, p, was set as 2 whereby only quadratic non-linearities of the system are captured and any higher order nonlinearities that might be present are ignored, and 2) system memory, L, was set as 60 msec. These model parameters were selected *a priori* given the compromise between model complexity and practical considerations of computational and experimental burdens.

**Figure 1.**
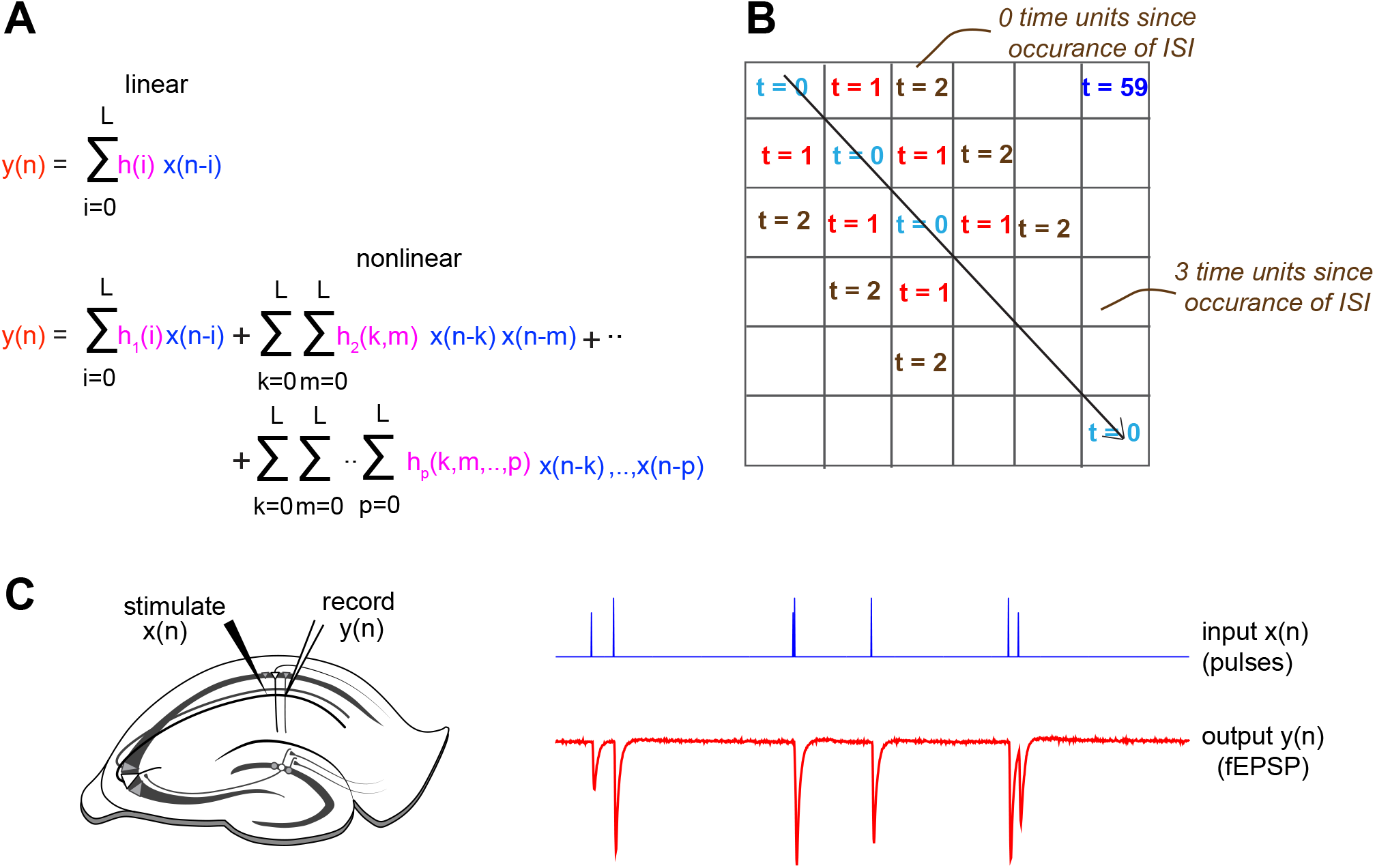
Using Volterra series expansion to identify the CA1 apical dendritic operations. **A,** Corresponding methods to fully characterize linear and nonlinear systems. *h(i)* fully characterizes a time-invariant linear system, and *h(i)* up to *hp (k,m,.,p)* fully characterize a nonlinear system with p^th^-order nonlinearities. **B,** *h(k,m)* is a 2-d symmetric matrix of weights. *h(k,m)* is a function of two parameters, the inter-spike-interval (ISI, or τ) represented by a fixed diagonal, and how far in the past the ISI occurred (moving along the diagonal). **C,** Schematic of slice electrophysiology stimulation and recording set up for the apical dendrites. Inputs *x(n)* were delivered to the Schaffer collaterals and outputs *y(n)* were recorded from the CA1 apical dendrites.

The term *h(i)*, the first order kernel, is a vector of weights on current and past single input values that contribute to the current output. The term *h(k,m)*, the second-order kernel, is represented by a symmetric matrix of weights describing the relative influence of past pairs of inputs on a current fEPSP (**Fig. 1B**)). The values on its diagonal slices indicate how the product of two input values τ time points apart affect a subsequent response occurring after various delays. Values further along a given diagonal (a fixed interval between inputs) reflect weights for the multiplication of neighboring inputs that occurred even further in the past relative to the current output (i.e., the memory of the system, and more simply, the CA1 *record* of its input history, **Fig. 1B**). A simplified schematic description for the pertinent variables of a second order Volterra series is illustrated in **Supplementary Material Figure 1**.

To estimate the first and second order kernels, we stimulated the Schaffer collaterals in young adult mouse hippocampal slices with pulse train *x(n)* and recorded the output signals *y(n)* from the proximal apical dendrites of the CA1 (**Fig. 1C**). For each slice (10 slices, n=20 sessions), input signals were delivered in two 15-minute sessions. Intervals between stimulation pulses were drawn from a Poisson distribution with a pulse rate of 2/sec and with amplitudes chosen uniformly from two values: *x_1_* which induced a sub-maximal fEPSP response *y_1_* of around 3-4 mV, and *x_2_* as a fraction of *x_1_* (either ½ or ¾). These input properties and experiment durations were carefully chosen through simulations to ensure kernel estimation accuracy (**Supplementary Figure 2**).

#### CA1 apical dendritic kernel estimates

Estimation of kernels is reformulated as a least-squares regression problem (see methods). Group kernel estimates were obtained using the last 12 minutes of each 15-minute session from each animal and after normalizing the input-output data. The linear (1^st^ order) kernel is an estimate of the filtering properties and reflects how the current and past input values are weighted to contribute to the generation of the fEPSP. Such linear weights reached a maximum in 10-15 msec and then exponentially decayed towards zero during the following 50 msec (**Fig. 2B**). The 2^nd^ order kernel showed that maximum nonlinear influence on a current response also occurred when past pulses were temporally close to each other (**Fig. 2C**). This is evident in the matrix plot, from the higher magnitude kernel values in slices immediately off the main diagonal. Second order main diagonal entries reflect the weights for the contribution of the squared values of the input in generating the response (**Fig. 2D**). The unit impulse response, which is the sum of the first order kernel and second order main diagonal entries, is by definition a system’s expected response to a delta-function (i.e., a stimulation pulse), which in the present case is the fEPSP (**Fig. 2E**). The result matched the complete waveform of a typical fEPSP recorded in the proximal apical dendrites (**Fig. 2A**). Note: individual animal *h(i), h(k,m)* and *h(k,k)* estimates are included in **Supplementary Figures 13-15**, respectively.

**Figure 2.**
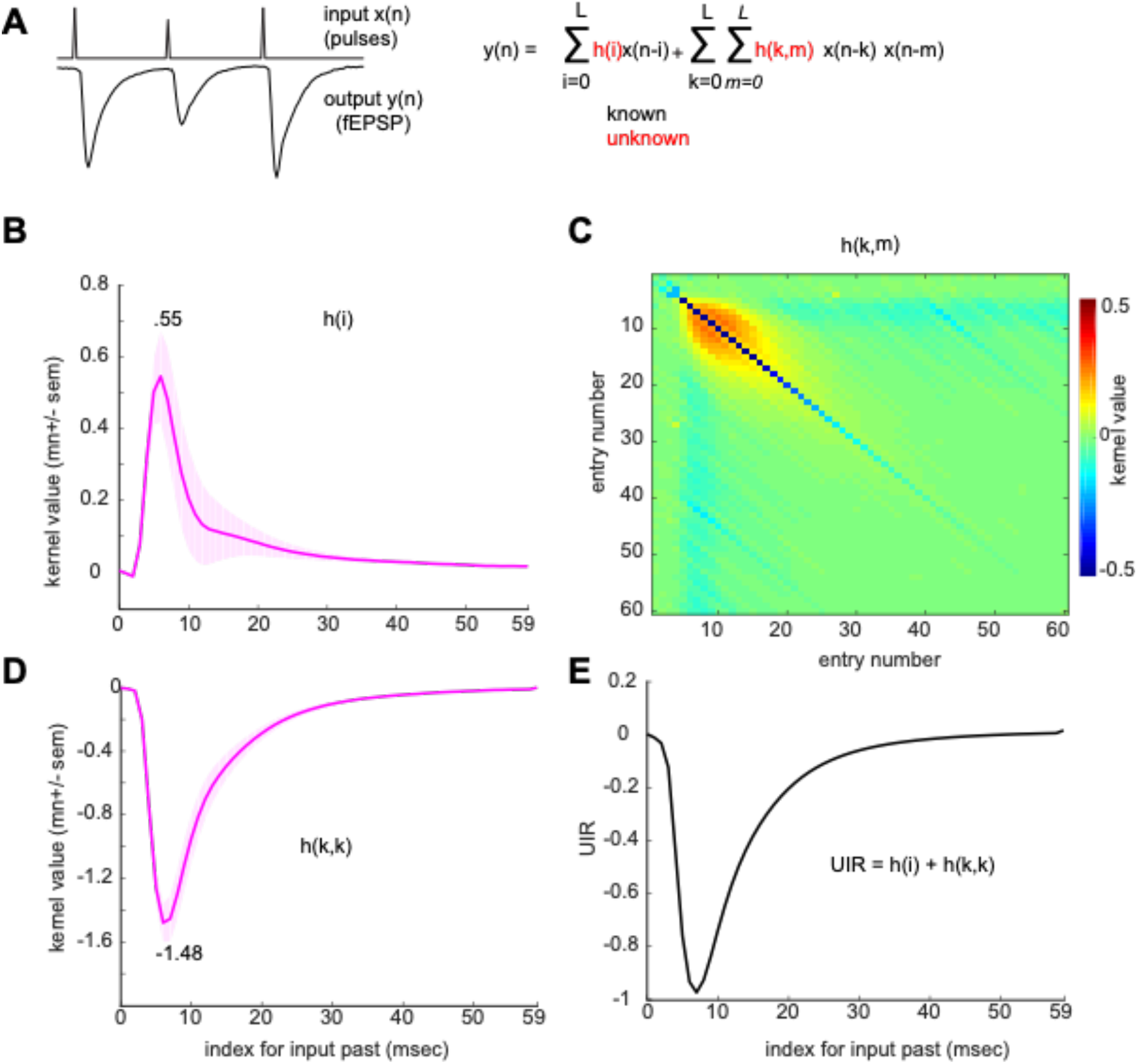
CA1 apical dendritic kernel estimates. **A,** Example input stimulation and output CA1 apical dendritic recording data (left) and second order truncated Volterra series expansion (right). **B-C,** the identified CA1 apical dendritic system; *h(i)* **(B)**, *h(k,m)* **(B)**. **D,** *h(k,m)* main diagonal slice, *h(k,k)*. **E,** system unit impulse response (UIR). Standard error shades reflects standard error of the mean (SEM) across sessions.

#### Derived kernels generalize across animals with high prediction accuracy

We then tested if the transfer function accurately reproduces outputs from slices from different animals that were not included in the original derivation. To do so, data from 9 animals were used to estimate the transfer function, which was then used to predict the CA1 output for the 10^th^ animal (Leave-one-out (LOO) cross validation, **Fig. 3A**). Normalized external prediction accuracy (normalization was relative to the magnitude of the true output; detailed in methods) across the 10 animals was 94.73%. The transfer function was therefore generalizable to new data, and reliably described the output across complex input patterns (**Fig. 3B**).

**Figure 3.**
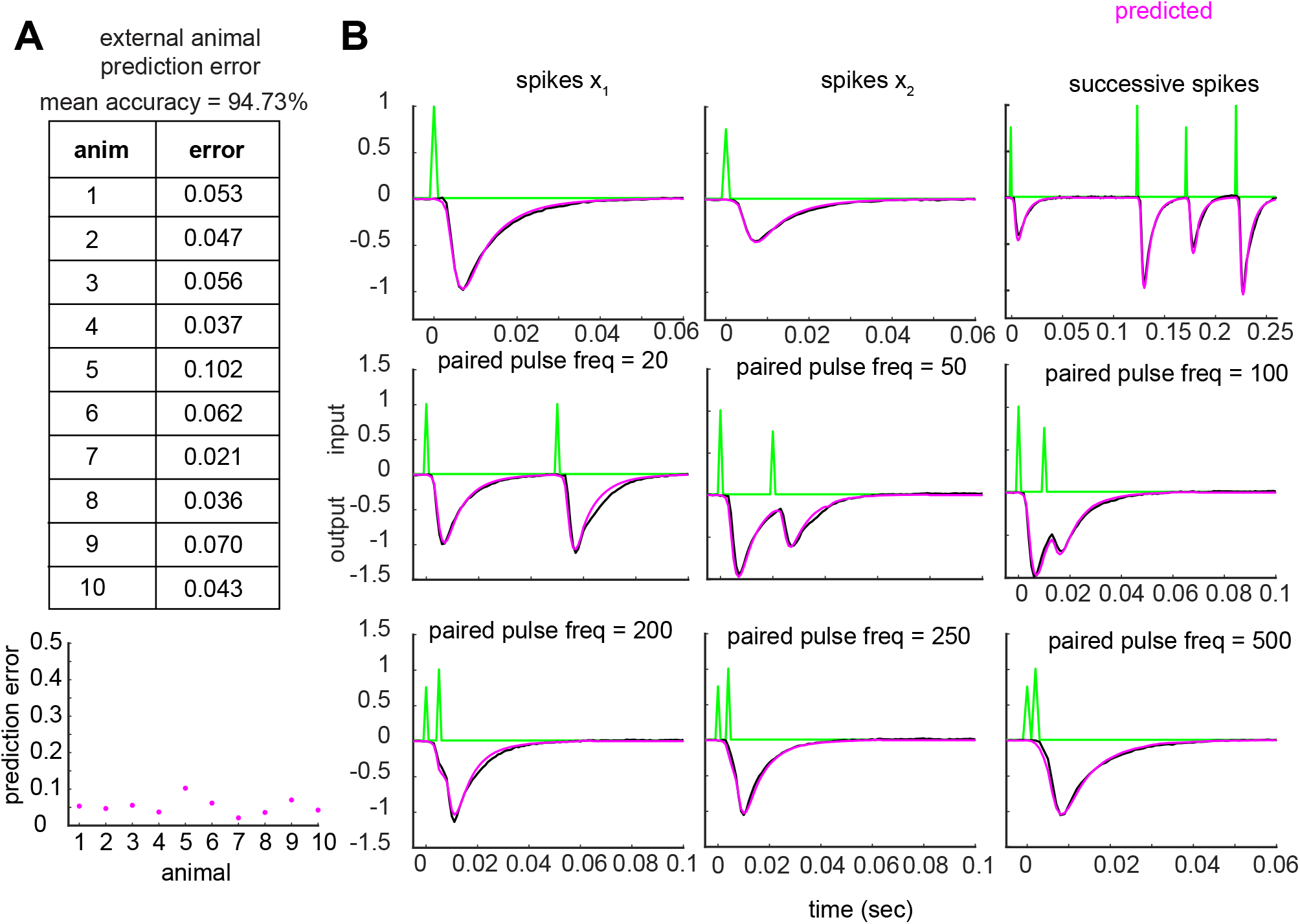
Estimated CA1 transfer function reliably predicts CA1 output on new animals. **A,** Table (top) and plot (bottom) displaying external animal prediction error (model generated using n-1 animals, with the test animal excluded). **B,** Example of predictions from animal 8 data for isolated spikes (top row, first two panels), successive spikes (top row, third panel), and paired pulses of variable inter-spike-intervals ISI (remaining). All examples were extracted from a continuous 15-min recording.

#### The nonlinear component of the apical dendritic operations outweighs the linear one in generating the dendritic response

The above findings raise questions about the relative contributions of the linear vs. nonlinear components of the equation to the fEPSP waveform. The magnitude of the 2^nd^ order kernel diagonal values (1.48) was larger than that of the 1^st^ order kernel (0.55, see **Fig. 2B,D**, above). Using these two values, the contribution of the linear component *(h(i))* and nonlinear component *(h(k,m))* to the peak output can be calculated as a function of input strength (number of afferent axons stimulated). The ratio of nonlinear-to-linear contribution to the response increases with increasing magnitudes of the input (the square of the input is larger, thereby applying *h(k,m)* weights to this larger square contributes more greatly to the output). For the CA1 apical dendritic system, the nonlinear component outweighs the linear one in generating the output at input values larger than ~0.4 (translates to 40% of the input that induces a submaximal response, **Fig. 4A**); see **Supplementary Figure 19** for individual animal estimates.

**Figure 4.**
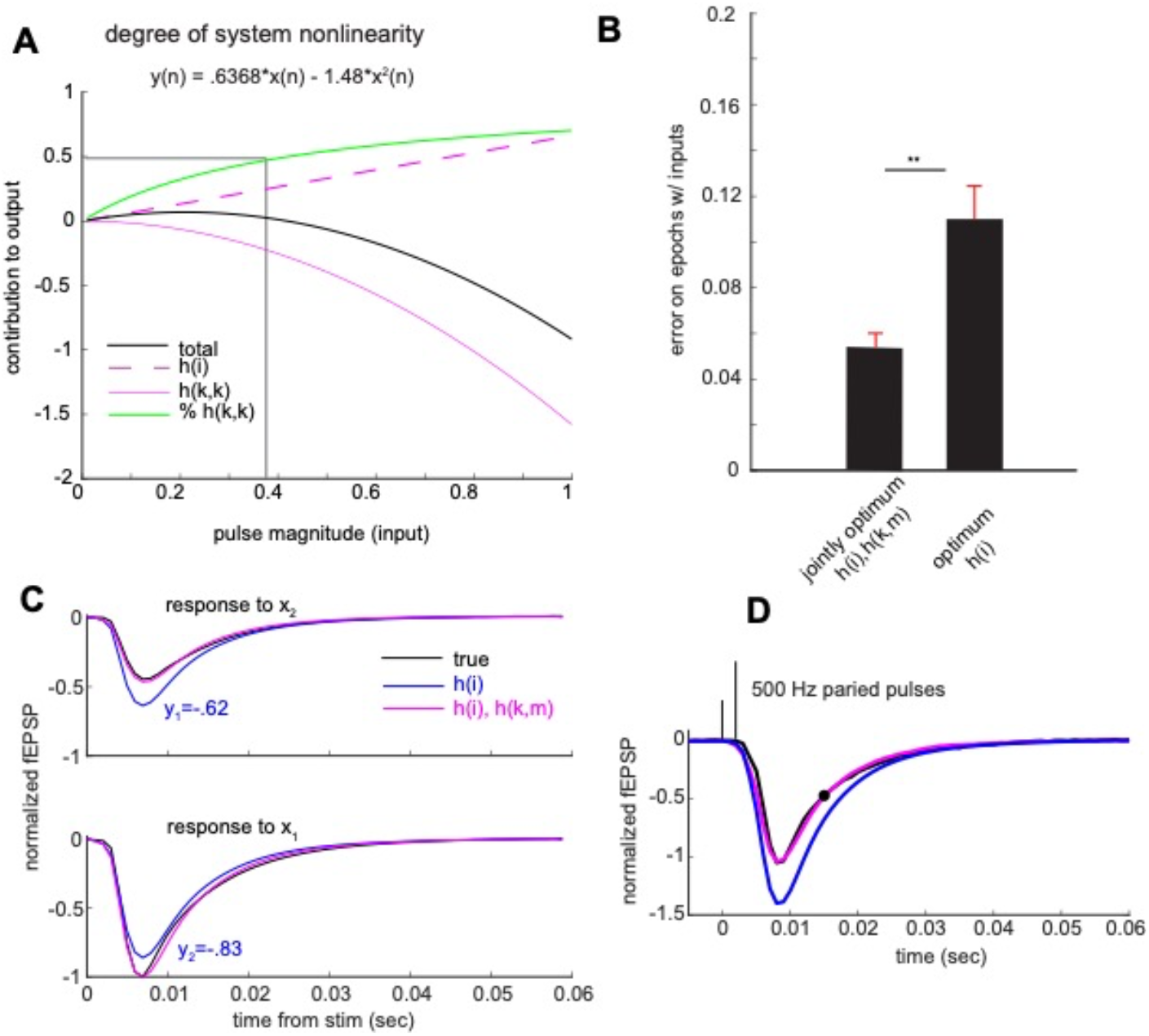
The nonlinear component of the apical dendritic operations outweighs the linear one in generating the dendritic response. **A,** Quantification of system nonlinearity. Using the peak values of the *h(i)* and *h(k,k)* (indicated in Fig. 2B,D), the predicted system output peak value (black) is plotted as a function of the input. Here, system output (black) is defined as a univariate value (the peak value of the fEPSP) rather than the continuous fEPSP waveform. The contributions of *h(i)* and *h(k,m)* to the output are displayed in magenta dashed and solid lines, respectively. The percentage of output generated by the nonlinear system component is shown in green. The intersection of the vertical and horizontal lines indicates that the nonlinear contribution to the output outweighs the linear one for input ranges larger than ~0.4. **B,** Differences in left-out animal prediction error using jointly optimum *h(i), h(k,m)* and the optimum *h(i)*, reflecting nonlinear and linear systems, respectively *(p* < 0.001, paired non-parametric permutation testing). **C,** Black traces reflect mean true response to x_1_ (top) and x_2_ (bottom) to isolated spike inputs (not preceded nor followed by spikes in the 1-sec and 0.25 sec periods relative to stimulation, respectively). Predicted waveforms by linear and nonlinear CA1 models are shown in blue and magenta, respectively. Linear CA1 model applies superposition (y2 is the same linear combination of y1 as x_2_ is of x1; y2 = ¾ * y1). **D,** An input of two spikes that are 2 msec apart is shown in black. Part of the predicted output value at 15ms (black circle) is generated by the nonlinear term whereby the product of the two input pulses is weighted by an entry of the matrix slice that is 2-off the diagonal.

The above results indicate that the nonlinear component of the transfer function is essential for generating the dendritic output. We quantified this by measuring the drop in output waveform prediction accuracy when using only a first order kernel compared to using both the first and second order kernels. To do so, we estimated two transfer functions; an optimum stand-alone *h(i)* kernel and jointly optimum *h(i)* and *h(k,m)* kernels. We found that prediction error on left-out data was significantly lower when using the transfer function of a nonlinear CA1 system in contrast to a purely linear system (*p* < 0.001, paired non-parametric permutation testing) (**Fig. 4B**).

The above analyses included multiple instances in which a single stimulation pulse was not preceded by another pulse for 1000 msec. Examination of these cases led to an informative result: an estimated linear system does not accurately predict changes in fEPSP amplitude associated with changes in single pulse stimulation current (**Fig. 4C**). Thus, nonlinearity is introduced by shifts in the number of active afferent fibers. This is of interest given the widely accepted argument that the CA1 system, in contrast to other hippocampal subdivisions, operates in a linear manner [13, 14]. The combination of different input sizes arriving in the recent past, in addition to their variable timing (pattern) which influences how the dendrites respond *(h(k,m))*, together lead to further error accumulation and thus further reduction in the predictive power of the 1^st^ order kernel with regard to the fEPSP waveform (**Fig. 4D** for an example). Including the estimated nonlinear transfer function allows for tracking of isolated inputs of variable magnitudes as well as arbitrary pulse patterns by accounting for the lag between all possible pairs of inputs and the time of occurrence of such lags.

#### New timing rules in the apical dendrites

The matrix of second order kernel weights provides a means for identifying essentially any 2^nd^ order nonlinear interactions between past inputs and present responses, many of which would go undetected in conventional physiological studies. To facilitate this application, we developed a pruned second order kernel which included values that were either below or above the 60^th^ and 80^th^ percentiles, respectively, of the mean divided by the standard deviation of entries across all sessions (consistently large negative and positive weights, respectively, across sessions) (**Fig. 5A**). As a general point, the matrix shows that consecutive pulses occurring in rapid succession (~2-11 msec) in the recent past (~5-40 msec) will depress current responses (less negative fEPSP; red pixels) while twin pulses separated by longer intervals (~20-59 msec) will exert a facilitative effect starting at 10 msec onward after delivery of the second pulse (blue pixels). Importantly, the initial segment of the slope of the second pulse (~1-10 msec) is largely unaffected by nonlinear components of the dendrites (green pixels). Given the high accuracy of predictions from the transfer function, we can be confident that these physiological rules describe outcomes when, as *in vivo*, target cells receive continuous input. We tested for the presence of these rules in the recordings, and this proved to be the case (**Fig. 5B-D)**.

**Figure 5.**
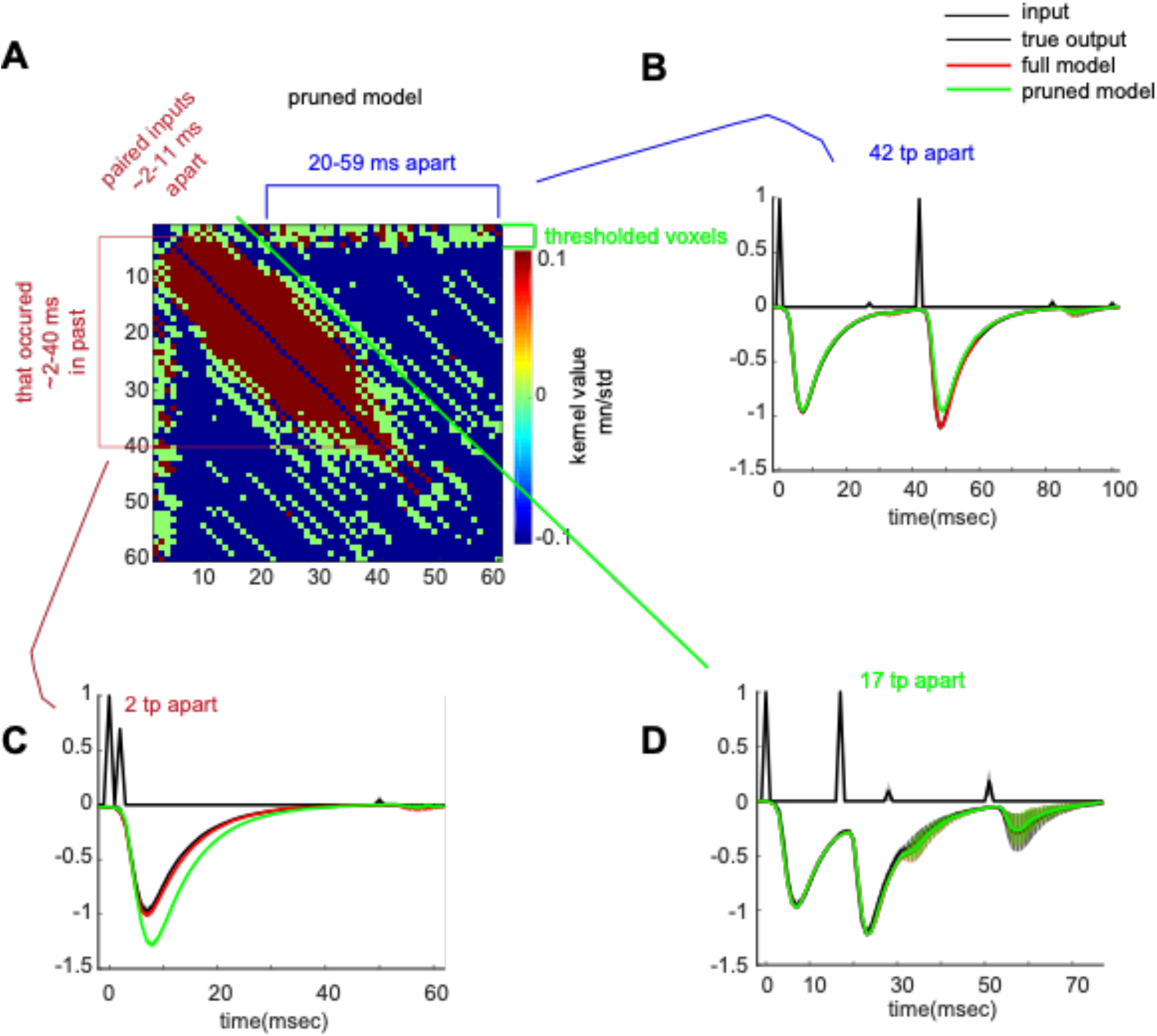
New timing rules in the apical dendrites. **A,** Pruned *h(k,m)* displays a thresholded matrix where included pixels are those in the top (red) and bottom (blue) 20% and 40^th^ percentiles, respectively, of the standardized mean (mean/standard deviation) kernel value distribution. Removed entries are indicated in green (such entries are close to 0 and thus the system does not employ them for output generation). Positive weights (red bracket) are applied for input pairs with an ISI of 2-11 msec. Negative weights are present for ISIs of 20-59 msec (blue pixels). **B,** Display of the importance of negative weights. Another pruned model was generated that excluded negative weights. Shown are full model predictions (with negative weights, including those for ISI of 42; red trace), pruned model (excludes negative weights, including those for ISI of 42; green trace), and true input and output (black trace). Negative weights are needed for an accurate prediction. **C,** same as in **(b)** but for positive weights. Positive weights are needed for an accurate prediction. **D,** same as in **(b)** but for green (thresholded) weights. Green pixels are not necessary for an accurate prediction.

### Signal transformation in CA1 basal dendrites

#### A transfer function for the CA3 projections to CA1 basal dendrites

Little is known about the dynamics of synaptic transmission in the CA1 basal dendrites. We accordingly employed the same system identification procedures used in the analysis of the apical dendrites to derive a transfer function for the basal domain. Stimulation pulses were delivered to a site in the distal stratum oriens of field CA1c (proximal CA1) approximately 400 μm removed from a recording pipette located at the same level of the lamina (**Fig. 6A**). As above, pulse intervals were taken from a Poisson distribution with mean spike rate of 2 Hz and pulse amplitude was drawn uniformly from two values x_1_ and a fraction of x_1_ (n=10 slices, n=37 sessions; **Table 2**). The 2^nd^ order Volterra series expansion was estimated as described in the earlier section on the apical dendrites. The obtained solution involved a sharp upward going first-order kernel *h(i)* which reflects system linear dynamics and an *h(k,m)* matrix which reflects system 2^nd^ order nonlinearities (**Fig. 6B-C**). The main diagonal slice from the matrix produced downward going weights over time, reflecting the weights for the contribution of the squared values of the input in generating the response (**Supplementary Figure 3A**). The unit impulse response – which is the system’s response to a single pulse – again matched the mean waveform of the basal fEPSP (**Supplementary Figure 3B**). Individual animal *h(i), h(k,m)* and *h(k,k)* estimates are included in **Supplementary Figures 16-18**, respectively.

**Figure 6.**
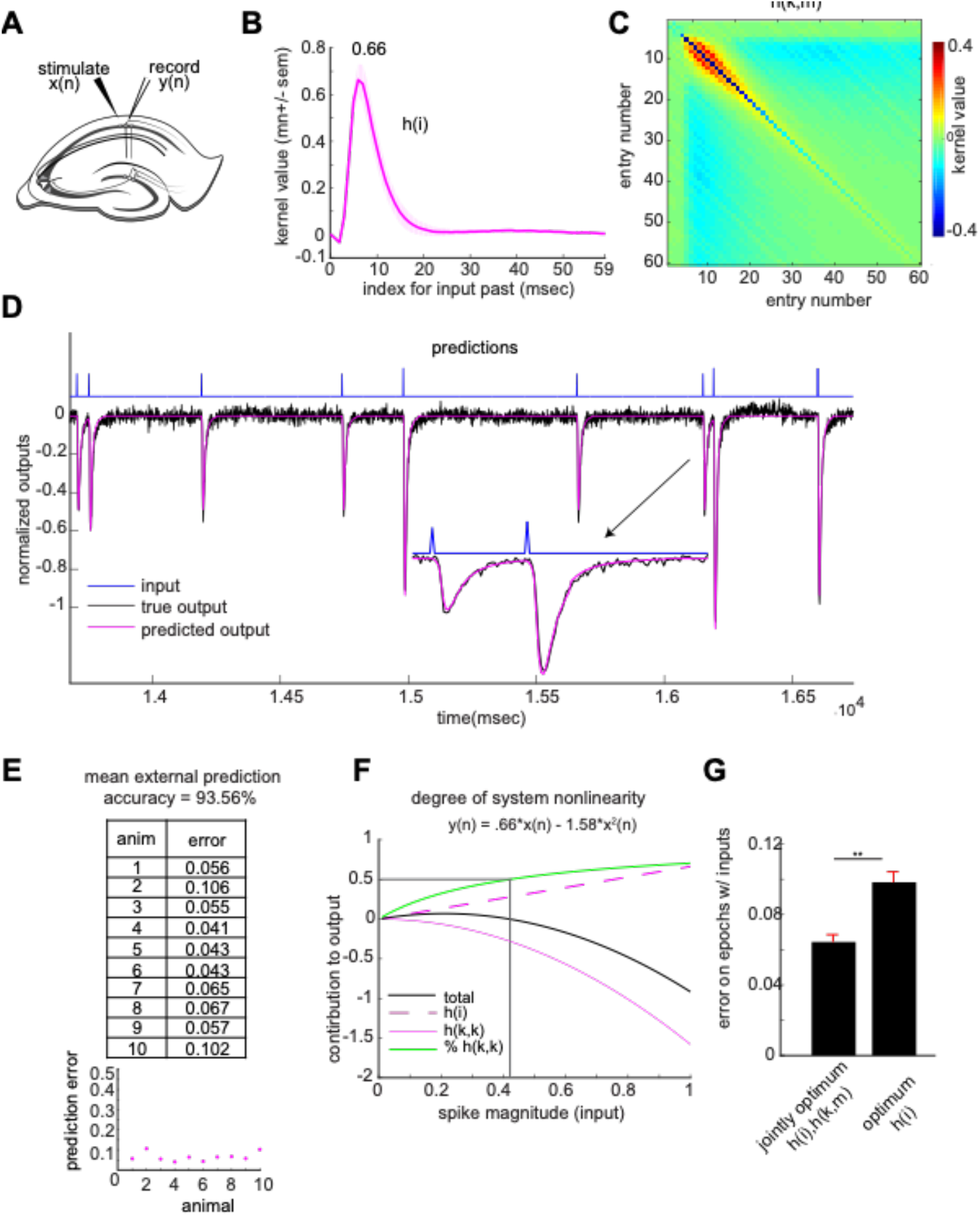
Signal transformation in the basal dendrites of CA1. **A,** Schematic of slice electrophysiology stimulation and recording setup for the basal dendrites. Inputs *x(n)* were delivered to the Schaffer collaterals and outputs *y(n)* were recorded from the CA1 basal dendrites. **B-C,** the identified CA1 basal dendritic system; *h(i)* **(B)**, *h(k,m)* **(B)**. **D,** Example trace from one recording session displays inputs (blue), true induced output (black) and externally predicted output (magenta). Inset shows a larger display of the paired spike segment of the trace. **E,** Table (top) and plot (bottom) display external animal prediction error (model generated using n-1 animals, with the test animal excluded). **F,** Quantification of system nonlinearity. Using the peak values of the *h(i)* (indicated in B) and *h(k,k)* (indicated in Supplementary Figure 3A), the predicted output peak value (black) is plotted as a function of the input. The contributions of *h(i)* and *h(k,m)* to the output are displayed in magenta dashed and solid lines, respectively. Percent of output generated by the nonlinear system component is shown in green. The intersection of the vertical and horizontal lines indicates that the nonlinear contribution to the output outweighs the linear one for input ranges larger than ~0.41. **G,** Differences in external animal prediction error using jointly optimum *h(i), h(k,m)* and the optimum *h(i)*, reflecting nonlinear and linear systems, respectively *(p* < 0.01, paired non-parametric permutation testing).

**Table 1.** Apical dendritic recording information regarding each recording session.

| **Apicals recording table** |  |  |  |  |
| --- | --- | --- | --- | --- |
| Experiment number | Animal number | Slice number | Trial number | Fraction for $x_1$ |
| 1 | 1 | 1 | 1 | 0.5 |
| 2 | 1 | 1 | 2 | 0.5 |
| 3 | 2 | 1 | 1 | 0.5 |
| 4 | 3 | 1 | 2 | 0.5 |
| 5 | 3 | 1 | 1 | 0.5 |
| 6 | 4 | 1 | 1 | 0.75 |
| 7 | 4 | 1 | 2 | 0.75 |
| 8 | 5 | 1 | 1 | 0.75 |
| 9 | 5 | 1 | 2 | 0.75 |
| 10 | 6 | 1 | 1 | 0.75 |
| 11 | 6 | 1 | 2 | 0.75 |
| 12 | 7 | 1 | 1 | 0.75 |
| 13 | 7 | 1 | 2 | 0.75 |
| 14 | 8 | 1 | 1 | 0.75 |
| 15 | 8 | 1 | 2 | 0.75 |
| 16 | 9 | 1 | 1 | 0.75 |
| 17 | 9 | 1 | 2 | 0.75 |
| 18 | 9 | 1 | 3 | 0.75 |
| 19 | 10 | 1 | 1 | 0.75 |
| 20 | 10 | 1 | 2 | 0.75 |

**Table 2.**
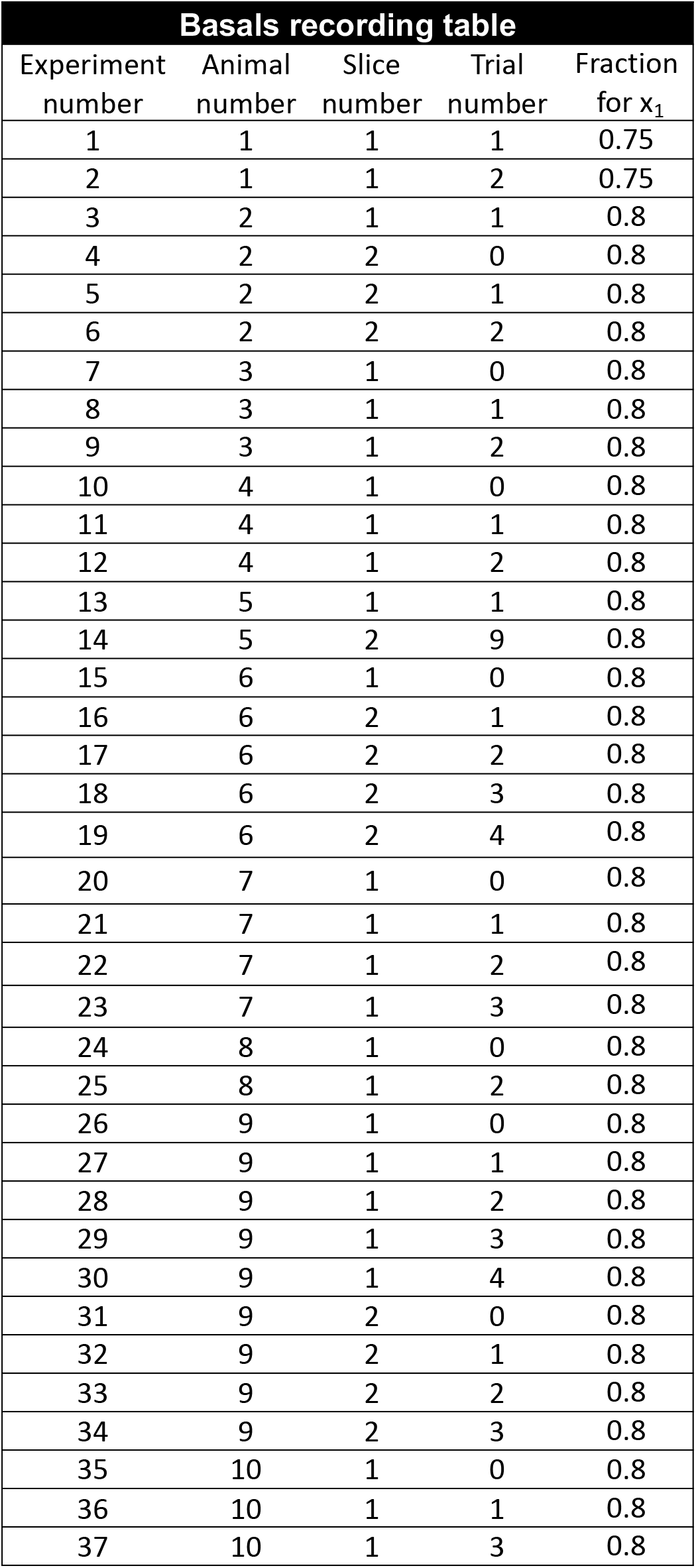
Basal dendritic recording information regarding each recording session.

| Basals recording table |  |  |  |  |
| --- | --- | --- | --- | --- |
| Experiment number | Animal number | Slice number | Trial number | Fraction for $x_1$ |
| 1 | 1 | 1 | 1 | 0.75 |
| 2 | 1 | 1 | 2 | 0.75 |
| 3 | 2 | 1 | 1 | 0.8 |
| 4 | 2 | 2 | 0 | 0.8 |
| 5 | 2 | 2 | 1 | 0.8 |
| 6 | 2 | 2 | 2 | 0.8 |
| 7 | 3 | 1 | 0 | 0.8 |
| 8 | 3 | 1 | 1 | 0.8 |
| 9 | 3 | 1 | 2 | 0.8 |
| 10 | 4 | 1 | 0 | 0.8 |
| 11 | 4 | 1 | 1 | 0.8 |
| 12 | 4 | 1 | 2 | 0.8 |
| 13 | 5 | 1 | 1 | 0.8 |
| 14 | 5 | 2 | 9 | 0.8 |
| 15 | 6 | 1 | 0 | 0.8 |
| 16 | 6 | 2 | 1 | 0.8 |
| 17 | 6 | 2 | 2 | 0.8 |
| 18 | 6 | 2 | 3 | 0.8 |
| 19 | 6 | 2 | 4 | 0.8 |
| 20 | 7 | 1 | 0 | 0.8 |
| 21 | 7 | 1 | 1 | 0.8 |
| 22 | 7 | 1 | 2 | 0.8 |
| 23 | 7 | 1 | 3 | 0.8 |
| 24 | 8 | 1 | 0 | 0.8 |
| 25 | 8 | 1 | 2 | 0.8 |
| 26 | 9 | 1 | 0 | 0.8 |
| 27 | 9 | 1 | 1 | 0.8 |
| 28 | 9 | 1 | 2 | 0.8 |
| 29 | 9 | 1 | 3 | 0.8 |
| 30 | 9 | 1 | 4 | 0.8 |
| 31 | 9 | 2 | 0 | 0.8 |
| 32 | 9 | 2 | 1 | 0.8 |
| 33 | 9 | 2 | 2 | 0.8 |
| 34 | 9 | 2 | 3 | 0.8 |
| 35 | 10 | 1 | 0 | 0.8 |
| 36 | 10 | 1 | 1 | 0.8 |
| 37 | 10 | 1 | 3 | 0.8 |

#### Derived kernels generalize across animals with high prediction accuracy

The derived transfer function reproduced fEPSPs across various stimulation patterns (**Fig. 6D)**. To quantitatively evaluate accuracy, the basal dendritic transfer function was estimated using data from n-1 animals and then used to predict the dendritic response on the nth animal (LOO). External prediction was done for all sessions and slices with the results summarized for each animal (**Fig. 6E**). The solution is shared from animal to animal, predicting the entirety of the synaptic response on a msec-by-msec basis with a mean external animal prediction accuracy of 93.56%. Calculations of the relative contributions of the linear and nonlinear elements yield a result similar to that for the apical dendrites: the nonlinear contribution outweighed the linear one as stimulation pulse intensity increased (**Fig. 6F**); see **Supplementary Figure 19** for individual animal estimates. As expected, the external prediction error significantly increases when estimating only an optimum *h(i)* solution to represent the optimum linear CA1 basal dendritic system (**Fig. 6G**)— evidence for the necessity of the nonlinear operations for reconstructing the dendritic response.

#### New signal transformation rules in the basal dendrites

A pruned version of the 2^nd^ order kernel was again obtained by including only entries whose mean divided by the standard deviation across slices was lower or higher than the 65^th^ and 87^th^ percentiles, respectively (**Fig. 7a**). With satisfactory agreement to the recorded data, the pruned matrix shows which input patterns generate more positive (red pixels) versus negative (blue pixels) fEPSPs by second order nonlinear dynamics, and that again, the slope response to the second pulse is largely unaffected by the nonlinear dynamics (green pixels) (**Fig. 7B-D** for examples).

**Figure 7.**
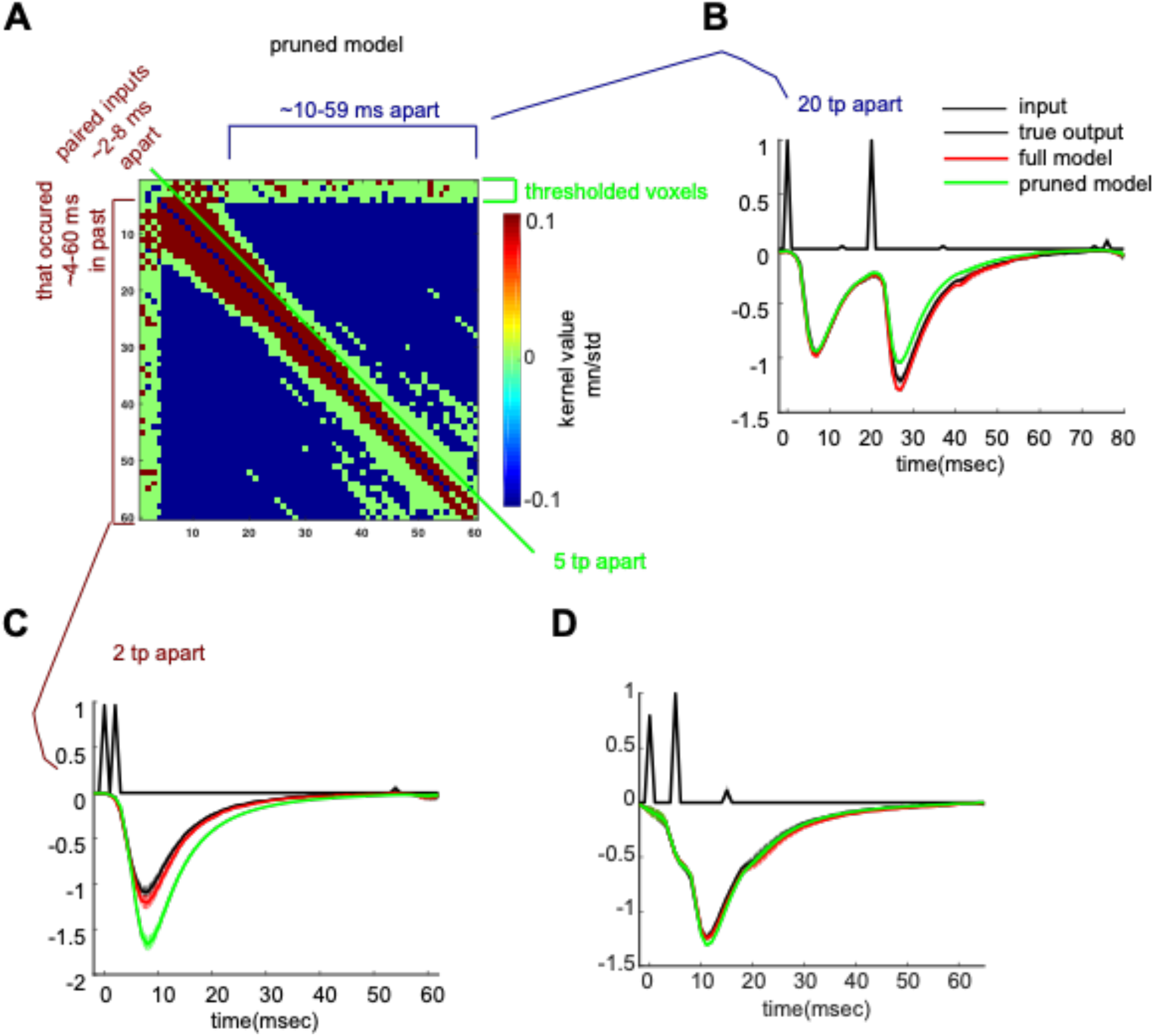
New timing rules in the basal dendrites. **A,** Pruned *h(k,m)* displays a thresholded matrix where entries in the top (red) and bottom (blue) 13^th^ and 65^th^ percentiles, respectively, of the standardized mean (mean/standard deviation) kernel value distribution were included. Removed entries are indicated in green (such entries are close to 0 and thus the system does not employ them for output generation). Positive weights (red bracket) are applied for input pairs with an ISI of 2-8 msec. Negative weights are present for ISI’s of 10-59 msec (blue pixels). **B,** Display of the importance of negative weights. Another pruned model was generated, whereby negative weights were excluded. Shown are full model predictions (includes negative weights, including those for the ISI of 20; red trace), pruned model (excludes negative weights, including those for the ISI of 20; green trace), and true input and output (black trace). Negative weights are needed for an accurate prediction. **C,** same as in **(b)** but for positive weights. Positive weights are needed for an accurate prediction. **D,** same as in **(b)** but for green (thresholded) weights. Green pixels are not necessary for an accurate prediction.

While the basal dendrites’ *h(k,m)* revealed nonlinear dynamics that were indeed present in the data, in examining paired pulse prediction errors, we discovered that *h(k,m)* predictions are inaccurate for some cases of paired pulses (**Supplementary Figures 4-7**). However, paired predictions held for most cases in the apical dendrites (**Supplementary Figures 8-11**). These observations indicate that the apical dendrites are sufficiently explained by 2^nd^ order dynamics, while the basal dendrites require a 3^rd^ order kernel or higher for complete characterization [15].

### Comparison of signal transformation in the apical vs. basal dendrites

#### Signal transformations in the apical differ from those in the basal dendrites

There were general similarities between the two dendritic domains; no significant differences were found between the basal and apical *h(i)* and *h(k,k)* (**Supplementary Figure 3 B-C**. Direct comparisons of *h(k,m)*, however, revealed significant differences in the two dendritic domains. For successive stimulation pulses whose ISI was 2-3 msec, the apical dendrites apply smaller weights on the fEPSP thus yielding a more negative fEPSP compared to the basal dendrites (blue pixels, **Fig. 8a**). The converse is true for a larger range of ISIs, anywhere from ~3-59 msec ISIs, whereby the apical dendrites apply larger weights, thereby generating less negative fEPSP decays compared to the basals (red pixels, **Fig. 8a**). To examine this further, diagonal slices of *h(k,m)* reflecting the weights for the input history of paired pulses arriving with an ISI of 30 msec apart were plotted separately (**Fig. 8B, top**). Such traces clearly revealed that compared to their basal counterparts, apical dendrites apply less negative weights (less facilitation) generating a less negative output ~7-29 msec after the occurrence of the second pulse (after the occurrence of the ISI event).

**Figure 8.**
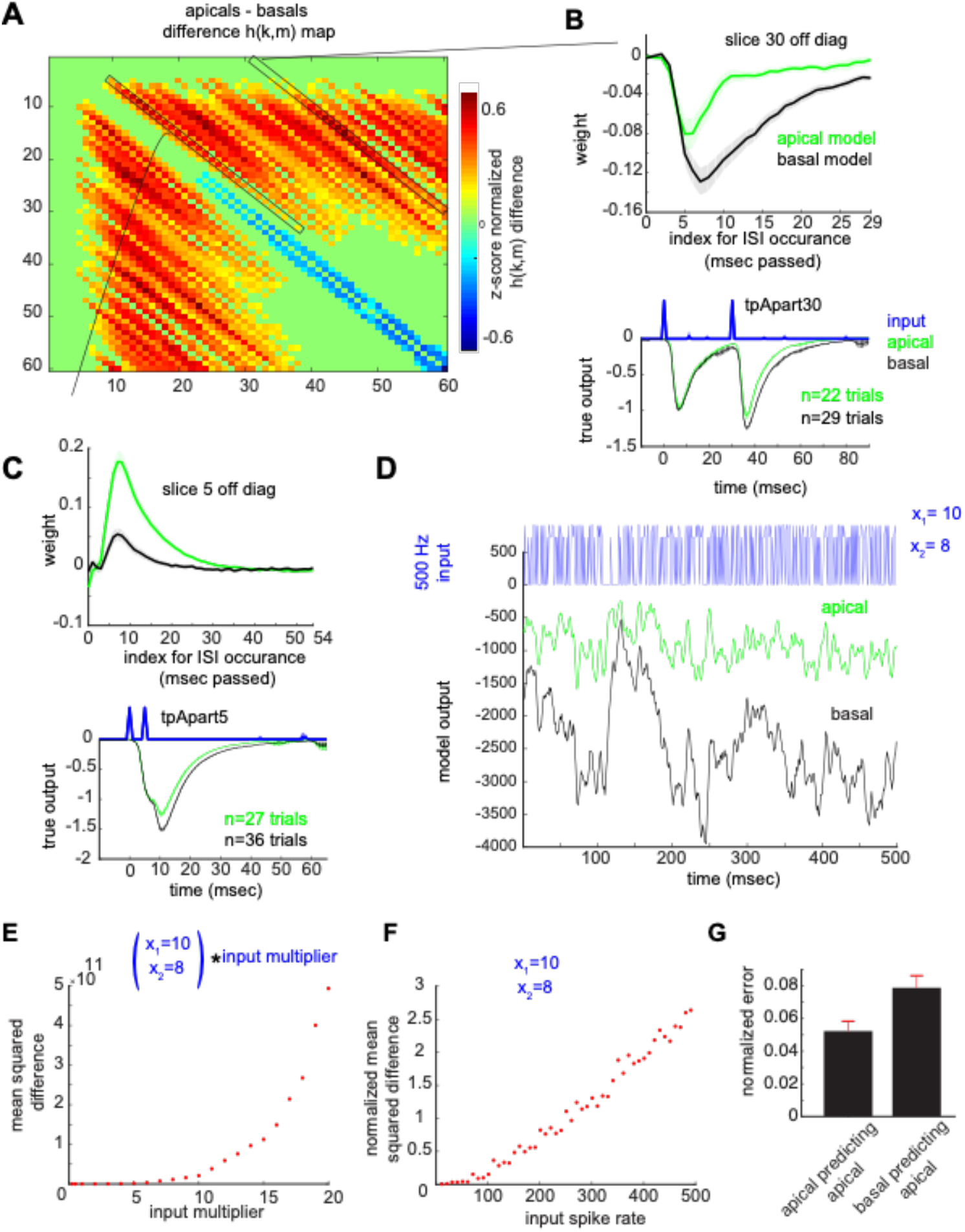
The apical transfer function cannot generate basal output signals and vice versa. **A,** z-score normalized apicals-basals difference *h(k,m)* map, cluster corrected for multiple comparisons (CBPT, p < 0.05, using null distribution cluster size to correct for multiple comparisons, and using sessions as observations). Green pixels are *h(k,m)* values that did not significantly differ between the two regions. Warm and cool pixels are significantly higher and lower *h(k,m)* values for the apicals compared to the basals, respectively. **B-C (top),** A group mean trace of kernel values across sessions for a given off diagonal slice in the matrix (slice 30 = 30 msec ISI (B, top) and slice 5 = 5 msec ISI (C, top) off the diagonal), indicated with black rectangles in A, for the apical (green) and basal (black) models. **B-C (bottom),** Mean input and output traces of all spikes with 30 msec (B, bottom) and 5 msec (C, bottom) ISIs pooled from all apical (green) and basal (black) recording sessions. Note the relatively less negative apical decay phase is consistent with the identified nonlinear system operations (B-C, top) whereby the addition of more largely weighted paired input products to the remaining terms in the VES yields a less negative output. **D,** A 500 msec segment of the apical and basal transfer function output to a Poisson input with mean rate of 500 Hz and spike magnitude uniformly drawn from x_1_ = 10 and x_2_ = 8. **E-F,** Mean squared difference between apical and basal transfer function outputs as a function of input magnitude (E) and input spike rate (F), respectively. **G,** External prediction error on pooled high amplitude successive spikes x_1_ and the 60 msec period following the second spike, for all spikes with ISI’s of 1 msec – 60 msec apart. Basal cross prediction error was significantly higher when predicting the output waveform in the apical dendrites, (*p* = 0.01, unpaired permutation testing, n = 100 permutations). Error shades and bars represent SEM across sessions.

We next confirmed the presence of such dynamics in the experimental data, using group mean outputs induced by ISIs of 30 msec (**Fig. 8B, bottom**). Indeed, the decay phase of the apical response to the second pulse, separated from the first by 30 msec, is less negative compared to that of the basals. Another example is shown for the *h(k,m)* slice reflecting weights for paired pulses with an ISI of 5 msec (**Fig. 8C, top**). The larger positive weights in the apicals compared to the basals should yield, again, a less negative fEPSP ~5-35 msec after the occurrence of the second pulse, which is indeed observed in the group average of such trial types (**Fig. 8C, bottom**). In addition to the significant differences identified in the wings of *h(k,m)*, it is also important to note that the magnitude of the raw (non-normalized) fEPSP differed between the domains; the basal domain generates a more compressed version than that of the apical domain (**Supplementary Figure 3E**). This suggests that the two domains’ *h(i)* and *h(k,k)* are comparably similar in waveform, but are rescaled versions of each other, while *h(k,m)* for *k* ≠ *m* differs significantly (**Supplementary Figure 3F**).

#### The apical transfer function cannot generate basal output signals and vice versa

Two characteristics of our data mask the extent to which the two dendritic domains are capable of generating different analog output waveforms. The first of these is the output magnitude, which is capped at ~1 due to the normalization. Yet, the contribution of the system nonlinearities to the output for both the apicals (**Fig. 4A**) and basals (**Fig. 6F**) increases with input magnitude, and it is in the nonlinear operations that the two systems differ. Secondly, our experimental protocol relied on a slow mean spike rate (2 Hz), meaning that on average, pairs of spikes were 500 msec apart. However, the weights in *h(k,m)* which significantly differed between the two systems cover a range of ISIs of 1-59 msec which translate to a spike frequency range of 17-1000 Hz. Therefore, our experimental protocol of small magnitude inputs and slow spike rates does not produce inputs that tap directly into the difference in dynamics between the two systems. We therefore took advantage of our discovery of such differences together with accurate transfer function estimates and examined the basal and apical model behaviors to inputs with increasing magnitudes and spike rates (**Fig. 8D**). Basal and apical model outputs are very different with high input magnitudes and spike rates. Quantifying the mean squared difference between the two system’s responses to the same input shows that the magnitude of the difference increases with increasing input magnitude **(Fig. 8E)** and spike rate **(Fig. 8F)**.

Lastly, as significant differences between the apical and basal dendritic transfer functions were observed at the level of the second order kernel *h(k,m)*, which reflects system operations on pairs of pulses, we tested cross-transfer function predictions on such successive pulses. We found a significant increase in external prediction error on pooled epochs with successive pulses (1-60 msec apart) from all animals, when using the basal transfer function to predict the output waveform to arbitrary inputs delivered in the apical dendrites (**Fig. 8G**).

### Identifying sources of prediction error

Several sources of prediction error were identified. First, CA1 dendritic system memory is longer than the 60 msec interval considered in the study. One way to demonstrate this is through the paired pulse analyses (**Supplementary Figures 4-11)**, where predicted traces deviate from true CA1 responses starting with the 60^th^ msec timepoint after the first pulse. After the 60^th^ msec, the predicted output response is generated by operations on the second pulse alone (the first pulse falls outside of the model’s 60 msec memory) while the CA1 response is still responding to both pulses (retained memory for the first pulse that occurred over 60 msec in the past).

Non-stationarity in the data is another source of prediction error (**Supplementary Figure 12**). While the majority of recordings appeared to be stationary (showed a constant induced response magnitude as a function of the recording durations, **Supplementary Figure 12A**), a subset was nonstationary (i.e., a gradual change in the magnitude, see **Supplementary Figure 12D**). Volterra kernels are only applicable for modeling stationary systems, as the kernels are assumed to be time invariant. They are estimated by minimizing the squared-error difference between the actual and predicted model outputs; therefore, for nonstationary data, the kernels will have higher prediction accuracy in the subset of the recording that is most similar in range to that of the group level induced CA1 responses (**Supplementary Figure 12F**).

A third source of error (more relevant for the apical model) was the variable *h(i)* across animals (see SEM shades in the grand average **Fig. 2B** and individual animal estimates **Supplementary Figure 13**). Variability in *h(i)* was visible in terms of both waveform shape and magnitude. This difference in *h(i)* is theoretically consequential and informs as to the nature of system output as a function of input (**Supplementary Figure 19 A-B**). For example, the *h(i)* estimate for apical animal 5 varied compared to estimates for other animals (**Supplementary Figure 13**). The magnitude of this *h(i)* coupled with animal 5’s *h(k,k)* estimate will dictate the linear and nonlinear contributions to the system output (as discussed for **Fig. 4A**). Because *h(i)* is large in magnitude, it will predominate in generating the output over a larger range of the normalized input (rightward shift in **Supplementary Figure 19C**). In addition, due to the fact that it is an upward going waveform, it will generate a *less negative* fEPSP for such input ranges. These properties were unique to this animal, hence the different input-output curve (**Supplementary Figure 19A**), and the highest external animal prediction error (**Fig. 3A**).

Lastly, examination of paired pulse prediction error suggests that a model with higher order than 2 is needed for the basal domain (this was clearly evident in **Supplementary Figure 7**, plots boxed in black).

## Discussion

The primary objective of the present work is to develop a systematic and concise mathematical characterization of signal transformation for inputs arriving from field CA3 and terminating in the dendrites of field CA1. This entailed analysis not only of inputs of different strength but more importantly the influences exerted by prior and current inputs on present responses. The endpoint measure was the generation of the entirety of the fEPSP which is a direct reflection of the primary unit (the excitatory post-synaptic current; EPSC) of communication between cortical regions. The system developed here predicted the size and waveform of the dendritic response, on a msec-by-msec basis, with a high degree of accuracy. Further reductions in prediction error will likely be obtained by extending basal system order beyond 2^nd^ order and dendritic system memory beyond 60 msec in accord with experimental results [16–18].

We found that deviations of CA1 system dynamics from linearity can be quantified in a generalizable manner and are dictated by the magnitude of the ratio of *h(k,m)* to *h(i)*. The larger the ratio, the earlier the deviation occurs at smaller values of the input. Nonlinearities made up more than 50% of output magnitude at normalized input values larger that ~0.4 (translates to 40% of the input which induces a submaximal response); at these values, the second-order kernel contributes more to the fEPSP waveform. This result does not accord with contemporary arguments that posit marked differences in non-linear input/output relationships for the dentate gyrus and field CA3 with CA1 having a more linear relationship [13, 14]. One effect of these proposed subfield differences is to allow for pattern separation by the divergent afferents of the dentate gyrus and pattern completion by the convergent interior of CA3 with CA1 providing reliable readout. There is empirical evidence for this general model, but our results suggest that adjustments are required for its output stage [13, 14, 19].

Field EPSPs are commonly used to evaluate the effects of experimental treatments such as drugs or genetic mutations on brain operations. Studies of this type typically employ widely spaced stimulation pulses and single measures of the evoked response such as slope or amplitude to test for changes from baseline; they only rarely include information about the dynamic properties of synapses as elicited by temporal patterns of inputs. The comprehensive and agnostic approach followed in the present studies has led to a readily implemented means for obtaining a far more complete description of synaptic operations. The *h(i*) and *h(k,m)* terms fully characterize the behavior of the 2^nd^ order apical system with a given memory and allow for a simple but accurate measurement of the distance in synaptic operations from a baseline state to that produced by a condition of interest. These features should result in a sizable gain in sensitivity and thus greater discrimination between the changes elicited by experimental manipulations.

Similar arguments apply to ongoing attempts to comprehensively describe the differences in signal processing by the various links in the hippocampal circuitry. Multiple studies have shown that the pertinent synapses respond in surprisingly different ways to repetitive stimulation delivered at frequencies corresponding to dominant EEG rhythms. For example, the apical branch of the CA3 to CA1 projections exhibits a conventional frequency facilitation effect to 5, 20, and 50 Hz trains [4] while the lateral perforant path connections to the dentate gyrus exhibit a form of low pass filtering [20]. These results are informative regarding differences in underlying neurobiological mechanisms but constitute special cases that do not lend themselves to the extraction of general rules of the type discussed above. We accordingly established a transfer function for the input to the basal dendrites and found it to be clearly different than that for the apical tree. The values for *h(k,m)* in the two domains differed because of evident discrepancies in the forward contributions of past events. Moreover, a limited number of cases in which the basal dendritic transfer function did not fully predict the effects of closely spaced pulses strongly suggest that a 3^rd^ or higher order kernel will be needed to fully characterize responses in this lamina. While techniques exist for estimation of larger order models while reducing the need for unrealistic amounts of experimental data, it is unknown *a priori* whether the functionals used in these approaches can represent the underlying ground truth Volterra kernels (see for example a discussion on the Laguerre Expansion Technique [12]). Clearly, a combination of modeling and experimental work will be needed to develop a conceptually robust, higher order description for the basal dendrites.

There are several possible contributors to the differences in signal processing by the apical vs. basal dendrites. We observed a larger variability in individual animal *h(i)* estimates in the apical compared to the basal domain (see standard error shades Fig. 2D vs. 2C). Since *h(i)* reflects the linear dynamics, that is for the dendritic system is largely passive conductance along the membrane, *h(i)* variability is likely a result of the more variable branching pattern of the apical dendritic membranes [21]. The small numbers of synapses activated by the weak stimulation pulses used in the present study are more likely to activate different electrotonic locations from experiment to experiment in the apical than basal dendrites. There were also differences between the two nodes in the non-linear aspects of the responses produced by multiple input pulses. For the dendritic system, nonlinearities are activated as thresholds for components in the synapse are reached. Relevant to this, pyramidal cells express a large number of diverse potassium and calcium channels, certain of which are not uniformly distributed along dendritic branches [22–24]. The channels affect the size and waveform of synaptic potentials, are sensitive to recent input history, and in many cases operate over time frames comparable to those in our stimulation protocol. Accordingly, regional differences in channel types [25] could account for much of the observed discrepancy between the 2^nd^ order kernels obtained for apical vs. basal domains. It is also the case that the two dendritic laminae contain different types of interneurons [11, 26, 27]. Interneuron subtypes vary substantially with regard to their activation requirements and firing patterns; moreover, there is evidence for selective targeting of particular combinations of GABAAR subunits [11]. There is thus a possibility that feedforward shunting inhibition elicited by repetitive afferent stimulation will not be the same in the basal dendrites as in their apical counterparts. Consistent with this are results showing that the GABA-mediated depression of the later EPSPs in a short sequence evoked by closely spaced stimulation pulses is much more pronounced in the str. radiatum than str. oriens [28].

The pyramidal neuron with distinct dendritic domains is a defining feature of mammalian cortex. There has been considerable interest in the possibility that its multiple dendritic domains allow for processing of input from diverse sources. We extend to this in showing that processing differs in these two domains. Given the presence of two distinct sets of computations on the same cell suggests that the production of maximal depolarization, and thus likelihood of evoked spiking, requires input patterns that lead to a net larger depolarization generated by the joint transfer functions. It is of interest in this regard that tests for relationships between *in vivo* spiking patterns in CA3 and CA1 have utilized 3^rd^ or higher order systems [17, 18]. In any event, the present findings are informative regarding hypotheses about the adaptive advantages of pyramidal cell architecture.

Finally, the findings reported here constitute a critical step in the development of hippocampal models from equations that incorporate a high degree of biological realism. Further progress in this direction will again require a transfer function for the conversion of EPSP sequences into spike patterns. Prior work using system identification techniques has established relationships between inputs and synchronous discharges of neuronal populations (‘population spikes’) [18, 29, 30] as well as dendritic shaft input to somatic membrane voltage output [31]. How these results might relate to the information-rich spike patterns used for communication across brain circuits is uncertain. Important work was also done whereby system identification on spike patterns was conducted using *in vivo* recordings from CA3 (input stage) and CA1 (output stage), although during well-defined behaviors which restricts the set of input patterns (those associated with the behavior) and thus cannot produce the generalized kernels obtained with sufficiently exciting inputs in an *in vitro* model. [17, 32, 33]. Moreover, correlations between the two subfields *in vivo* could in part reflect near simultaneous input to both regions from a third region such as the entorhinal cortex or septum, areas known to synchronize activity within the hippocampus. Nonetheless, the results provide special cases that must be satisfied by universal models. But assuming that reliable mathematical relationships for the EPSP-to-spike pattern transition can be developed, then it should be possible to assemble the diverse transfer functions for each of the links and nodes in the hippocampal circuit into a single predictive model. This would be a significant step towards the development of biologically realistic models of the brain.

## Acknowledgements and Funding

Funding for this project was provided by the National Institutes of Health grants R01MH115932 and R01AG053555 (to M.A.Y.), ONR N00014182114 (to G.S.L.) and training grant support T32NS45540 and T32GM008620 (to S.G.), T32MH119049 (to A.L.) and individual NRSA support (F30AG069406 to S.G.). We would like to thank Dr. Conor Cox, Dr. Ben Gunn, Prof. David Darmon and Prof. Paul Rapp for helpful discussions.

## Supplementary Data ^i^

## MATERIALS AND METHODS

### Animals

Experiments were conducted using 20 male mice (10 C57BL/6N) 2-3 months of age (**Tables 1-2**); 10 animals were utilized for apical and 10 for basal recordings. Animals were group-housed (4-5 per cage), on a 12-hr light/dark cycle and with food and water *ad libitum*. All procedures were approved by the UC Irvine Institutional Animal Care and Use Committee and in line with the NIH guidelines for the Care and Use of Laboratory Animals.

### Hippocampal slice electrophysiology

Animals were sacrificed between 10:00am-11:00am. Acute hippocampal slices were prepared as described previously [34]. Briefly, transverse slices of the middle third of the left hippocampus (360 um thickness) were cut using a McIlwain chopper and collected in chilled artificial CSF (ACSF) which contained in mM (124 NaCl, 3 KCl, 1.25 KH_2_PO_4_, 1.5 MgSO_4_, 26 NaHCO_3_, 2.5 CaCl_2_, and 10 dextrose). Slices were then transferred to the interface chamber kept at 31±1°C and had 60-70 mL/hr infusion of oxygenated ACSF. Slices were left for 1.5-2 hours to equilibrate prior to recording.

A stimulating electrode (twisted nichrome) was placed in the CA3-CA1 Schaffer collaterals and a glass recording electrode (2 M NaCl filled, 2-3 MΩ) was used to record extracellular field potentials from the CA1 dendrites. For apical dendritic recordings, the stimulating and recording electrodes were placed in the CA1b stratum radiatum with the stimulating electrode slightly offset towards the stratum pyramidale, and the recording electrode in the middle of the stratum radiatum [34]. In most animals, stimulation intensity (*x_1_*) was set to induce a ~3.5-4 mV population-spike free fEPSP. For basal dendritic recordings, the stimulating and recording electrodes were placed in the CA1b stratum oriens with the recording electrode in the middle of the stratum oriens and the stimulating electrode adjacent to it. In most animals, stimulation intensity (*x_1_*) was set to induce a ~1 mV population-spike free fEPSP.

### Recording protocol

For each animal, a 10-minute stable baseline session was recorded before experiment initiation. Baseline stimulation consisted of a single pulse every 20 seconds. After ensuring stable baseline responses, two or more 15-min sessions were recorded per animal. Slices were left to stabilize for 5 minutes without stimulation between the 15-min sessions. Stimulation patterns were random with spike times drawn from a Poisson distribution (lambda of 2 spikes/second) and amplitudes uniformly drawn from two values (*x_1_* and *x_2_*: *x_1_* induced a 3-4 mV and ~1 mV fEPSP in the apicals and basals, respectively and *x_2_* was a fraction of x_1_, **Table 1-2**). Recording duration and patterns were explored and determined through simulations such that they yielded accurate parameter estimates (**Supplementary Figure S2**).

### Equipment

The stimulation signal *x(n)* was generated digitally in MATLAB 2018a (The MathWorks, Natick, MA), converted to an analog signal using 16-channel data acquisition hardware (DA-16, A-M Systems), which was then connected to a stimulus isolation unit (2200, A-M Systems) enabling the delivery of current to the stimulating electrode. DA-16 was also used to digitally record the delivered analog input *x(t)* and CA1 analog output *y(t)*, sampled at 10,000 Hz and saved in MATLAB. Both stimulus delivery and recording implemented through the DA-16 were interfaced using MATLAB and its Data Acquisition Toolbox for Measurement Computing.

### Analysis: Simulations

Prior to initiation of data collection, simulations were conducted to aid in the appropriate experimental design for accurate estimates of 2^nd^ order, 60-msec memory Volterra kernels. Ground truth first- and second-order Volterra kernels were generated, using knowledge of the structure of these kernels (symmetric second-order kernel) and candidate CA1 dynamics (**Supplementary Figure 2A**). The ground truth first-order kernel was defined as an fEPSP. This is because for a linear system, *h(i)* is the response to a delta or spike input and the CA1 dendritic field generates an fEPSP to a spike input. Using these ground truth kernels, we tested the nature of the input and amount of data needed for accurate kernel estimation.

First, we found that increased randomness in the input increases the likelihood of tapping into the second-order structure. This is because such random input would likely include all possible τ, or ISIs, and kernel weights for all possible τ would be utilized and reflected in the output. Then, such kernel weights can be recovered with the input/output data. We therefore decided to deliver inputs at random times pulled from a Poisson distribution, which also mimicked the underlying distribution of neural spike timing (**Supplementary Figure 2B**). The Poisson parameter lambda was set to 2 spikes/second. A slow spike rate was chosen to prevent overstimulating the slice thereby affecting its dynamics during the recording and because this spike rate was used in prior work [15, 35, 36].

Second, we found that a binary input leads to estimation bias due to multicollinearity – where the first-order kernel and the diagonal elements of the second-order kernel become coupled. This is because the first-order kernel provides weights for individual input values and the diagonal of the second-order kernel provides weights for the square of such values. With a binary input (0 and a non-zero value), the square of the input vector becomes a linear combination of it leading to multicollinearity and estimation bias. We therefore randomly and uniformly drew the spike amplitude from two values, x_1_ and x_2_ (x_1_ induced ~3-4 mV and ~1mV responses in the apicals and basals, respectively, and x_2_ was a fraction of x_1_, see **Tables 1-2**).

Finally, for the desired model order – a 60 msec kernel length and up to a second-order kernel, we quantified the normalized estimation error as a function of data duration (**Supplementary Figure 2C-D**). The error reached a plateau of 0 with 13-15 minutes for a single session.

### Analysis: Data curation and preprocessing

Data curation and preprocessing were done offline in MATLAB. The recorded input signal was used to identify the input timestamps. The detected timestamps were then used to remove stimulation artifacts from the CA1 record. Local points (17 data points ~1.7 ms; 3 time points (tp) before and 14 sample points after) around the artifact peak were replaced by propagating the CA1 signal immediately before the artifact using the slope of the signal at the boundaries of the artifact 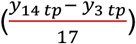 plus noise pulled from a normal distribution with 0 mean and the standard deviation of the CA1 artifact free record weighted by 1/16. Then, other large amplitude measurement noise data points were replaced by the mean of the stimulation artifact-free recording plus noise pulled from a normal distribution (0 mean and the standard deviation of the CA1 artifact free record). The data was then downsampled to 1000 Hz. All individual animal kernel estimates took place subsequently using this data (non-normalized).

For group analysis, additional preprocessing was implemented and consisted of within animal 1) pre-stimulation normalization and 2) normalization of fEPSP amplitude relative to the mean fEPSP induced by x_1_. Pre-stimulation normalization accounts for slow drifts in the data, which offset the starting value of the induced fEPSP. Each value following a stimulus was z-score normalized relative to the mean and standard deviation of a 100 msec (1000 data points, before downsampling) period prior to stimulation that does not overlap with a preceding CA1 response (after an 80 msec duration of a prior response) from a prior stimulus. For closely occurring successive stimulations, however, this baseline period did not exist. Therefore, successive stimuli occurring proximally in time shared the same pre-stimulus baseline period which occurred before the train.

After pre-stimulus baseline normalization, all values were normalized to the negative peak of the mean fEPSP trace induced by x1. This enabled all *y(n)* output values to be bounded between 0 and ~−1 for all animals. Additionally, this enabled normalization of the input signal across animals; whereby the x_1_ inducing ~ −1 normalized response was arbitrarily mapped to 1 for all animals. Then, x_2_ became simply equal to the original fraction used during the experiment – either 0.5, 0.75, or 0.8.

### Analysis: Kernel estimation

For each 15-min session recorded per animal, the last 12-min (most stationary subset) was utilized to obtain individual animal estimates for subsequent group analysis. This choice accounted for the tradeoff between stationarity and sufficient data for accurate estimation (**Supplementary Figures 2**). The session exclusion criteria included: 1) measurement noise preventing the stimulation detection, 2) non-invertibility of the design matrix, 3) recording error, 4) lack of slice stability (non-stabilizing fEPSP slope/peak values), and 5) high amplitude, temporally sustained, measurement noise.

The curated CA1 output signal was used as the *y(n)* vector, and an equal length spike vector, *x(n)*, was re-arranged into a matrix, *X_design_* such that the kernel estimation is reformulated into a least squares regression problem *y(n)* = *X_design_*\* *h_all_(n)* and solved using the pinv function in MATLAB. *h_all_(n)* contains all kernels (both first- and second-order) in vector format. It is important to note that in generating the design matrix, we took advantage of the symmetry of the second-order kernel and only estimated the diagonal and upper triangular elements of the kernel to reduce the number of unknown parameters.

### Analysis: normalized prediction error

Normalized prediction error was quantified as the sum of the squared error (predicted - true) normalized by the sum of the squared true output [29]. This measure was calculated for stimulation epochs (each stimulation time point and the following 60msec data). Such epochs are ones where non-zero predictions were made, otherwise the system is assumed to be at rest (predictions for system output equal 0).

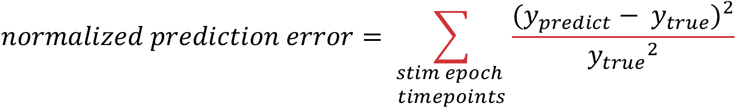

### Analysis: h(k,m) pruning

For each dendritic domain, a pruned version of *h(k,m)* was obtained. Pruning was done to remove coefficients that were near 0 and thereby obtain a sparser model that performs similarly to the full model. The pruning was done by examining the distributions of values for each entry in *h(k,m)* across sessions. Only entries with consistently large positive weights (large positive mean divided by standard deviation) and consistently large negative weights (large negative mean divided by the standard deviation) were kept, and the others removed. This was done by gradually increasing the percentile thresholds for mean/standard deviation upper and lower bound cutoffs. For each threshold, pruned model performance (prediction error) was tracked. The final pruned models (retained apical entries: lower than the 60^th^ and higher than the 80^th^ percentiles; retained basal entries: lower than the 65^th^ and higher than the 87^th^ percentiles) were chosen such that a high degree of pruning is achieved with no significant difference in prediction accuracy compared to the full model. For both dendritic domains, *h(k,m)* entry value distribution was skewed to the left (higher probability of negative weights). For this reason, the lower bound is as large as the 60-65^th^ percentile; values below these percentiles were negative, values between 60-65^th^ and 80-87^th^ were primarily 0, and values above the 80-87^th^ percentiles were positive for the apicals and basals, respectively.

### Analysis: paired pulse predictions

Paired pulse predictions were examined in each node to test whether the system is indeed second order (**Supplementary Figure 4-7** for basals and **Supplementary Figure 8-11** for basals). This was done by detecting from all recorded sessions all paired pulses with ISIs anywhere from 1 to 60 msec, irrespective of other pulses occurring in-between.

### Analysis: differences in kernel estimates between nodes

Significant testing for differences in *h(k,m)* kernel values between the apical and basal dendritic domains were identified using cluster-based permutation testing [37]. Briefly, this involved calculating a t-statistic in each entry, between the apical and basal *h(k,m)* 2-D observed matrices (n = 20 for the apicals, and n = 37 for the basals), thereby generating an *observed t-map*. The observed t-map was then compared to a null distribution of t-maps generated over 1000 unpaired permutations. In each permutation, condition labels were shuffled such that all *h(k,m)* matrices estimated from apical and basal recordings were randomly allocated to two fake groups with the same discrepancy in sample size (n = 20 for condition 1 and n = 37 for condition 2, hence unpaired permutations). For each entry, the observed t-value was compared to the null t-value in the same entry, in terms of the number of standard deviations the observed value was greater or less than the mean of the null t-value, thereby generating a z-map. The z-map was then converted to a *p*-map in which a *p*-value for each entry was obtained. To correct for multiple comparisons (number of entries tested), clusters of contiguous entries with *p* < 0.05 were identified and compared to the null-distribution cluster size. Observed clusters with sizes larger than the 95^th^ percentile of those from the null distribution were considered significant after correction for multiple comparisons. Significant testing for differences in *h(i)* between the two regions was conducted using the same procedure, but applied on a 1-D vector of entries rather than a 2-D matrix of entries [37].

### Analysis: Statistical analyses

Significance testing on prediction error was done using non-parametric paired permutation testing with 100 permutations. Significance testing on differences in true-versus-predicted paired pulse responses was done using the 1-D cluster-based permutation testing discussed in the prior section (‘*Analysis: differences in kernel estimates between nodes*’).

## Footnotes

i All supplementary data can be found on FigShare Project link below: https://figshare.com/proiects/Signal_Transformations_and_New_Timing_Rules_of_Hippocampal_CA3_to_CA1_Synapses/140107 Also accessible via DOI: 10.6084/m9.figshare.c.6014533

**Individual Figures and DOI links are:**

Supplementary Figure 19. https://doi.org/10.6084/m9.figshare.19878289.v1

Supplementary Figure 18. https://doi.org/10.6084/m9.figshare.19878286.v1

Supplementary Figure 17. https://doi.org/10.6084/m9.figshare.19878283.v1

Supplementary Figure 16. https://doi.org/10.6084/m9.figshare.19878277.v1

Supplementary Figure 15. https://doi.org/10.6084/m9.figshare.19878271.v1

Supplementary Figure 14. https://doi.org/10.6084/m9.figshare.19878265.v1

Supplementary Figure 13. https://doi.org/10.6084/m9.figshare.19878256.v1

Supplementary Figure 12. https://doi.org/10.6084/m9.figshare.19878259.v1

Supplementary Figure 11. https://doi.org/10.6084/m9.figshare.19878250.v1

Supplementary Figure 10. https://doi.org/10.6084/m9.figshare.19878241.v1

Supplementary Figure 9. https://doi.org/10.6084/m9.figshare.19878223.v1

Supplementary Figure 8. https://doi.org/10.6084/m9.figshare.19878214.v1

Supplementary Figure 7. https://doi.org/10.6084/m9.figshare.19878205.v1

Supplementary Figure 6. https://doi.org/10.6084/m9.figshare.19878193.v1

Supplementary Figure 5. https://doi.org/10.6084/m9.figshare.19878181.v1

Supplementary Figure 4. https://doi.org/10.6084/m9.figshare.19878154.v1

Supplementary Figure 3. https://doi.org/10.6084/m9.figshare.19878100.v1

Supplementary Figure 2. https://doi.org/10.6084/m9.figshare.19877206.v2

Supplementary Figure 1. https://doi.org/10.6084/m9.figshare.19853845.v2

## REFERENCES

1. Edwards, R.H., The neurotransmitter cycle and quantal size. Neuron, 2007. 55(6): p. 835–58.

2. Bellocchio, E.E., et al., The localization of the brain-specific inorganic phosphate transporter suggests a specific presynaptic role in glutamatergic transmission. J Neurosci, 1998. 18(21): p. 8648–59.

3. Dittman, J.S. and T.A. Ryan, The control of release probability at nerve terminals. Nat Rev Neurosci, 2019. 20(3): p. 177–186.

4. Jackman, S.L., et al., The calcium sensor synaptotagmin 7 is required for synaptic facilitation. Nature, 2016. 529(7584): p. 88–91.

5. Miki, T., et al., Actin-and Myosin-Dependent Vesicle Loading of Presynaptic Docking Sites Prior to Exocytosis. Neuron, 2016. 91(4): p. 808–823.

6. Miki, T., et al., Two-component latency distributions indicate two-step vesicular release at simple glutamatergic synapses. Nat Commun, 2018. 9(1): p. 3943.

7. Wang, K., et al., Distinct Ca2+ sources in dendritic spines of hippocampal CA1 neurons couple to SK and Kv4 channels. Neuron, 2014. 81(2): p. 379–87.

8. Zamponi, G.W. and K.P. Currie, Regulation of Ca(V)2 calcium channels by G protein coupled receptors. Biochim Biophys Acta, 2013. 1828(7): p. 1629–43.

9. Katz, B. and R. Miledi, The role of calcium in neuromuscular facilitation. J Physiol, 1968. 195(2): p. 481–92.

10. Jackman, S.L. and W.G. Regehr, The Mechanisms and Functions of Synaptic Facilitation. Neuron, 2017. 94(3): p. 447–464.

11. Schulz, J.M., et al., Dendrite-targeting interneurons control synaptic NMDA-receptor activation via nonlinear alpha5-GABAA receptors. Nat Commun, 2018. 9(1): p. 3576.

12. Marmarelis, V., Nonlinear dynamic modeling of physiological systems. 2004: Wiley-IEEE Press.

13. Knierim, J.J. and J.P. Neunuebel, Tracking the flow of hippocampal computation: Pattern separation, pattern completion, and attractor dynamics. Neurobiol Learn Mem, 2016. 129: p. 38–49.

14. Yassa, M.A. and C.E. Stark, Pattern separation in the hippocampus. Trends Neurosci, 2011. 34(10): p. 515–25.

15. Sclabassi, R.J., et al., Nonlinear systems analysis of the hippocampal perforant path-dentate projection. I. Theoretical and interpretational considerations. J Neurophysiol, 1988. 60(3): p. 1066–76.

16. Koutsoumpa, A. and C. Papatheodoropoulos, Short-term dynamics of input and output of CA1 network greatly differ between the dorsal and ventral rat hippocampus. BMC Neurosci, 2019. 20(1): p. 35.

17. Song, D., et al., Nonlinear dynamic modeling of spike train transformations for hippocampal-cortical prostheses. IEEE Trans Biomed Eng, 2007. 54(6 Pt 1): p. 1053–66.

18. Gholmieh, G., et al., An efficient method for studying short-term plasticity with random impulse train stimuli. J Neurosci Methods, 2002. 121(2): p. 111–27.

19. Guzowski, J.F., J.J. Knierim, and E.I. Moser, Ensemble dynamics of hippocampal regions CA3 and CA1. Neuron, 2004. 44(4): p. 581–4.

20. Amani, M., et al., Rapid Aging in the Perforant Path Projections to the Rodent Dentate Gyrus. J Neurosci, 2021. 41(10): p. 2301–2312.

21. Mihaljevic, B., et al., Comparing basal dendrite branches in human and mouse hippocampal CA1 pyramidal neurons with Bayesian networks. Sci Rep, 2020. 10(1): p. 18592.

22. Migliore, R., et al., The physiological variability of channel density in hippocampal CA1 pyramidal cells and interneurons explored using a unified data-driven modeling workflow. PLoS Comput Biol, 2018. 14(9): p. e1006423.

23. Ariav, G., A. Polsky, and J. Schiller, Submillisecond precision of the input-output transformation function mediated by fast sodium dendritic spikes in basal dendrites of CA1 pyramidal neurons. J Neurosci, 2003. 23(21): p. 7750–8.

24. Johnston, D., et al., Dendritic potassium channels in hippocampal pyramidal neurons. J Physiol, 2000. 525 Pt 1: p. 75–81.

25. Kirizs, T., et al., Distinct axo-somato-dendritic distributions of three potassium channels in CA1 hippocampal pyramidal cells. Eur J Neurosci, 2014. 39(11): p. 1771–83.

26. Klausberger, T. and P. Somogyi, Neuronal diversity and temporal dynamics: the unity of hippocampal circuit operations. Science, 2008. 321(5885): p. 53–7.

27. Cossart, R., et al., Interneurons targeting similar layers receive synaptic inputs with similar kinetics. Hippocampus, 2006. 16(4): p. 408–20.

28. Arai, A., J. Black, and G. Lynch, Origins of the variations in long-term potentiation between synapses in the basal versus apical dendrites of hippocampal neurons. Hippocampus, 1994. 4(1): p. 1–9.

29. Dimoka, A., et al., Modeling the nonlinear properties of the in vitro hippocampal perforant path-dentate system using multielectrode array technology. IEEE Trans Biomed Eng, 2008. 55(2 Pt 1): p. 693–702.

30. Berger, T.W., et al., Restoring lost cognitive function. IEEE Eng Med Biol Mag, 2005. 24(5): p. 30–44.

31. Cook, E.P., et al., Dendrite-to-soma input/output function of continuous time-varying signals in hippocampal CA1 pyramidal neurons. J Neurophysiol, 2007. 98(5): p. 2943–55.

32. Berger, T.W., et al., A hippocampal cognitive prosthesis: multi-input, multi-output nonlinear modeling and VLSI implementation. IEEE Trans Neural Syst Rehabil Eng, 2012. 20(2): p. 198–211.

33. Song, D., et al., Sparse Large-Scale Nonlinear Dynamical Modeling of Human Hippocampus for Memory Prostheses. IEEE Trans Neural Syst Rehabil Eng, 2018. 26(2): p. 272–280.

34. Wang, W., et al., Memory-Related Synaptic Plasticity Is Sexually Dimorphic in Rodent Hippocampus. J Neurosci, 2018. 38(37): p. 7935–7951.

35. Berger, T.W., et al., Nonlinear systems analysis of the hippocampal perforant path-dentate projection. II. Effects of random impulse train stimulation. J Neurophysiol, 1988. 60(3): p. 1077–94.

36. Berger, T.W., et al., Nonlinear systems analysis of the hippocampal perforant path-dentate projection. III. Comparison of random train and paired impulse stimulation. J Neurophysiol, 1988. 60(3): p. 1095–109.

37. Cohen, M.X., Analyzing neural time series data: theory and practice. 2014.

