## Supplementary Figures for "Signal Transformations and New Timing Rules of Hippocampal CA3 to CA1 Synapses"

Gattas et al.

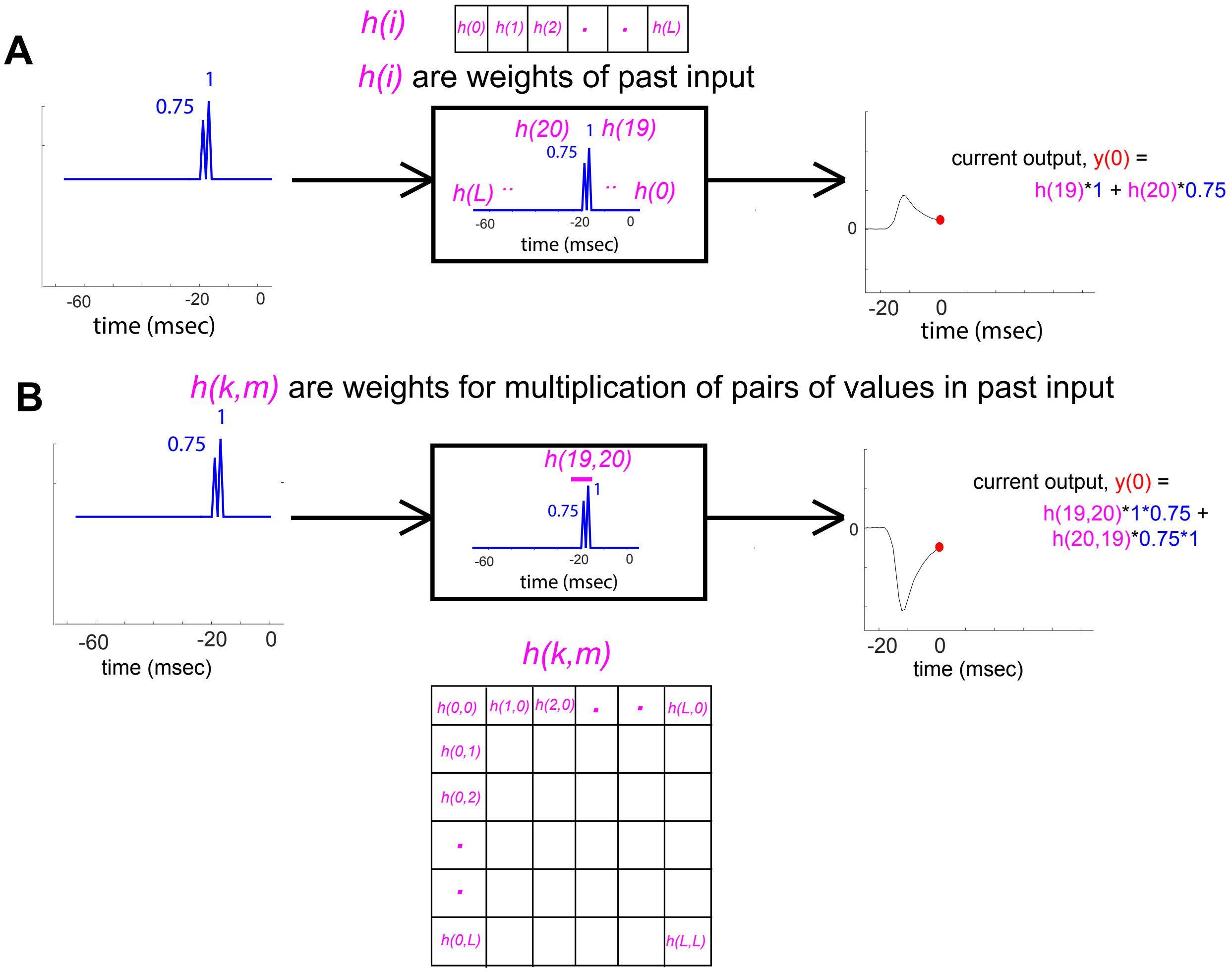


**Supplementary Figure 1. Nonlinear system identification**

**A-B,** Schematic representation of how *h(i)* (A) and *h(k,m)* (B) reflect system linear and second-order nonlinear operations, respectively. Blue traces preceding the arrow are inputs, black traces following the arrow are outputs and the blocks represent the system. System operations are reflected by the kernels *h(i)* and *h(k,m)* which are applied on incoming inputs to yield instantaneous output values. **A,** The term *h(i)*, the first-order kernel, is a vector of weights applied on current and past input values (up to L time-points in the input record) that contribute to the current output value (red circle). **B,** The term *h(k,m)*, the second-order kernel, is represented by a symmetric matrix; its τ's diagonal slices reflect how the output is influenced by products of neighboring input values τ timepoints apart occurring up to L samples in the past. Note: Output traces in A and B are not exact, but approximations to serve as an illustration.

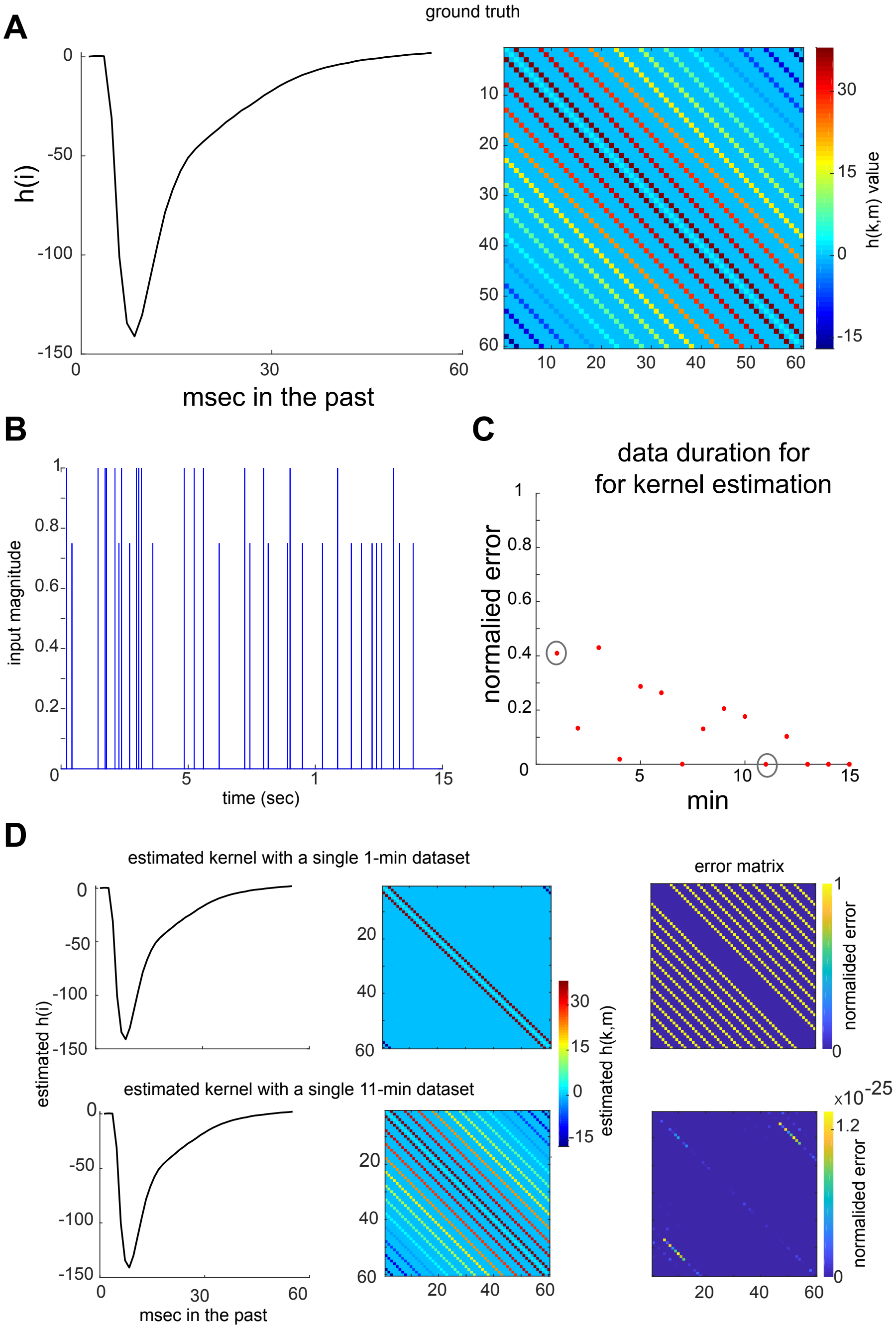


**Supplementary Figure 2. Simulations to identify recording duration and input type needed for accurate and unbiased kernel estimation.**

**A,** Artificially generated ground truth first (left) and second (right) order kernels. **B,** example of nonbinary ($x_1$ =1 or $x_2$ = 0.75* $X_1$) input with ISI from a Poisson distribution and spike amplitude from a uniform distribution. **C,** Normalized estimation error as a function of data duration. **D,** Example kernel estimates (left) and *h(k,m)* error matrix (right) using 1 minute (top) and 11 min (bottom) data, whose total errors are circled in **C.**

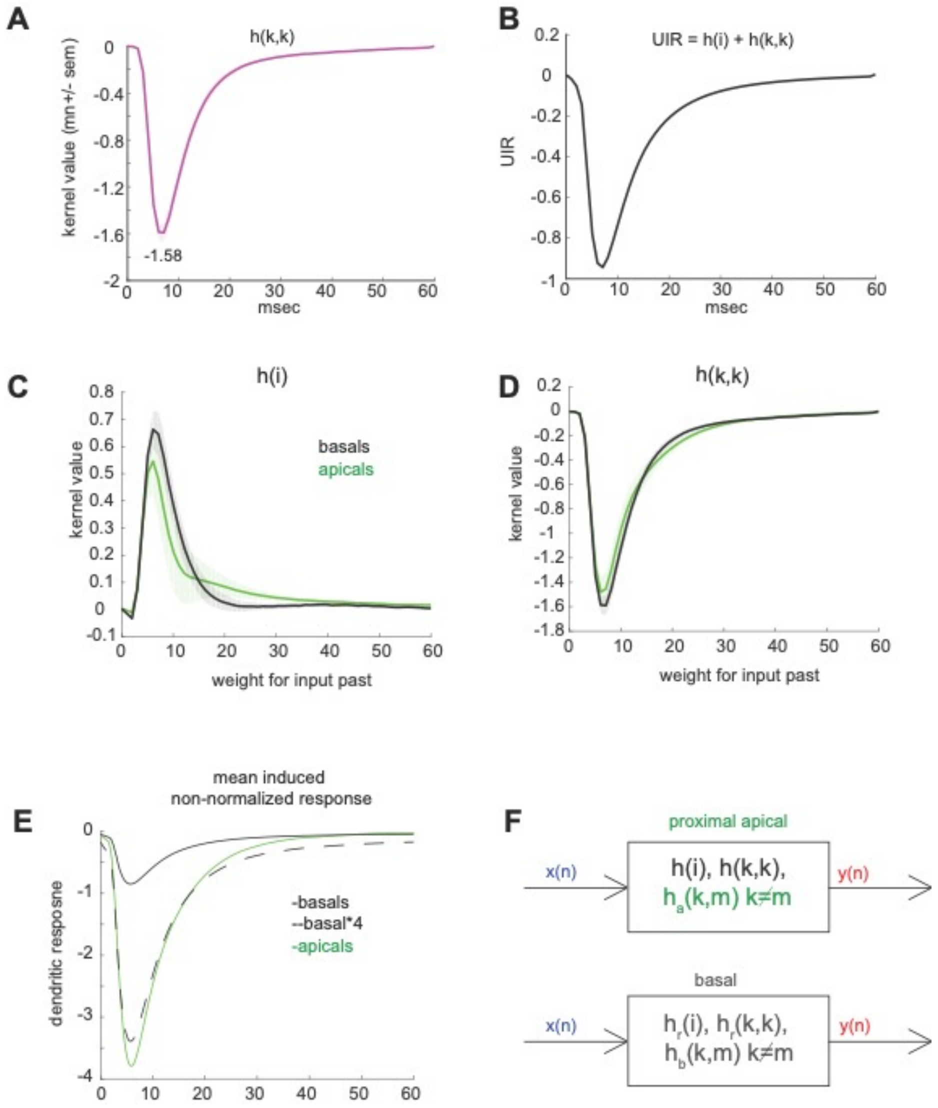


**Supplementary Figure 3. Aspects of the basal dendritic system transfer function and its relation to the apical system.**

**A,** Session mean *h(k,m)* main diagonal slice, *h(k,k)*. **B,** Session mean system unit impulse response (UIR). **C-D,** Session mean **(C)** *h(i)* and **(D)** *h(k,k)* estimates for the apical and basal systems (n = 20 and n = 37 sessions pooled from 10 animals per region, respectively). No significant *h(i)* nor *h(k,k)* differences were observed between the two dendritic domains (cluster-based permutation testing (CBPT), $p > 0.05$). **E,** Group mean induced response $y_1$ to all high amplitude input pulses $x_1$ aligned to the time of the input pulse and averaged for apical (green) and basal (black) recordings (n = 14,049 trials and n = 26,308 trials pooled from all sessions per region, respectively). Note the differences in the scaling of the raw induced fEPSP (before magnitude normalization). Dashed black line is a basal waveform rescaled by a factor of 4. **F,** Block diagram representation of transfer function for the proximal apical (top) and basal (bottom) systems. $h_r(i)$ and $h_r(k,k)$ are rescaled versions of the same kernel waveforms for the apicals *h(i)* and *h(k,k)*, yielding a compressed basal fEPSP. $h_a(k,m)$ and $h_b(k,m)$ are the offdiagonal weights of the apical and basal *h(k,m)* nonlinear operations, respectively, which are known to differ (**Fig. 8A**). Error shades represent standard error of the mean across sessions.

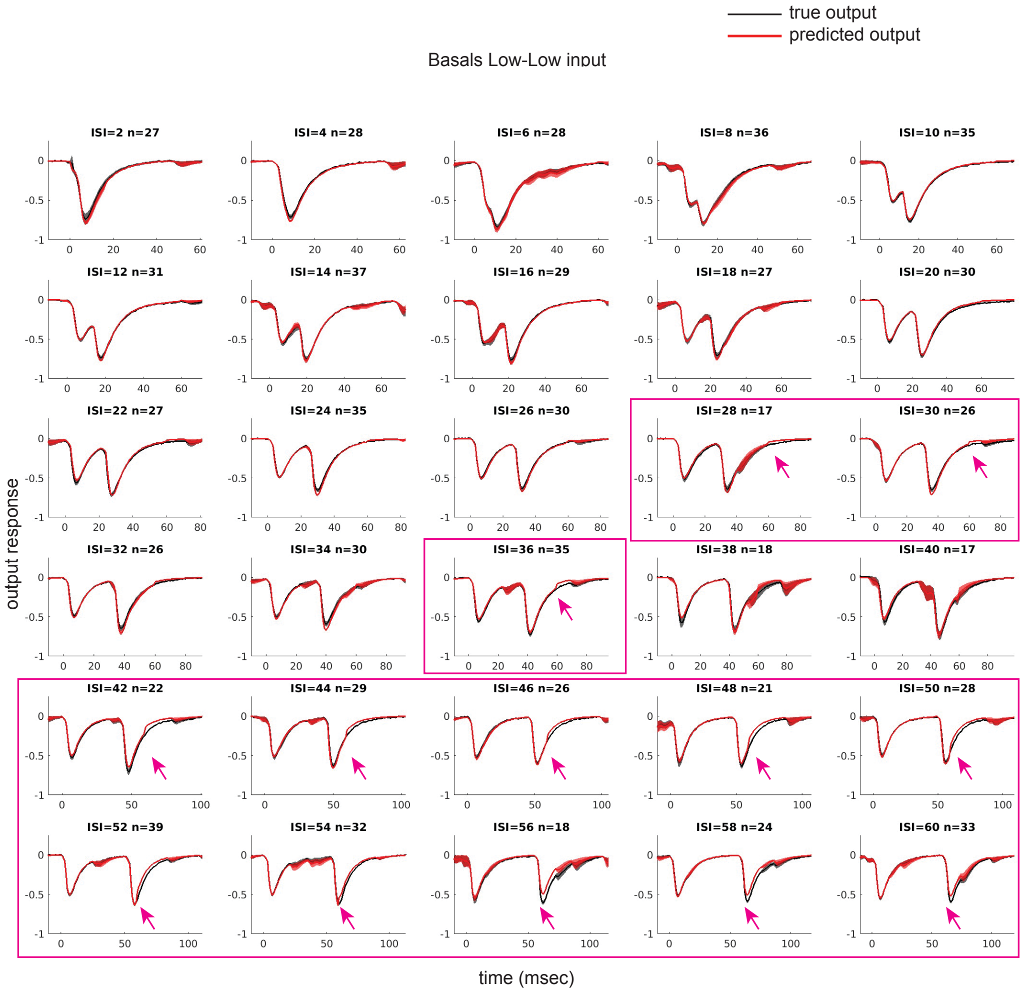


**Supplementary Figure 4. Paired pulse predictions for the basal dendrites (low magnitude for both pulses)**

True (black) and predicted (red) basal dendritic responses for paired pulse inputs with ISIs ranging from 2-60 msec in steps of 2 msec. For each plot, the ISI and number of trials used to generate the mean traces are indicated. Cluster based permutation testing (CBPT) was performed to identify time segments that significantly differed between the true and predicted conditions (1000 permutations). CBPT was only performed on the 60-msec time period after the first pulse since predicted time points after such periods are the model's response to the second pulse treating it as an isolated pulse (model has no memory for first pulse). ISIs for which significant differences were found are indicated by black boxed plots and the time duration that significantly differed between the true and predicted traces is indicated with a green line. No significant differences were found between the true and predicted responses for all ISIs plotted. Magenta boxed plots are a demonstration that system memory is larger that 60 msec. In those examples, the model's predicted traces deviate from the CA1's true response starting at the 60$^{th}$ msec onward, since at this point, the predicted output response is generated by operations on the second pulse alone while the CA1's response is still responding to both pulses (retained memory for the first pulse that occurred over 59 msec in the past). SEM reflects standard error of the mean across pooled trials with a given ISI from all sessions.

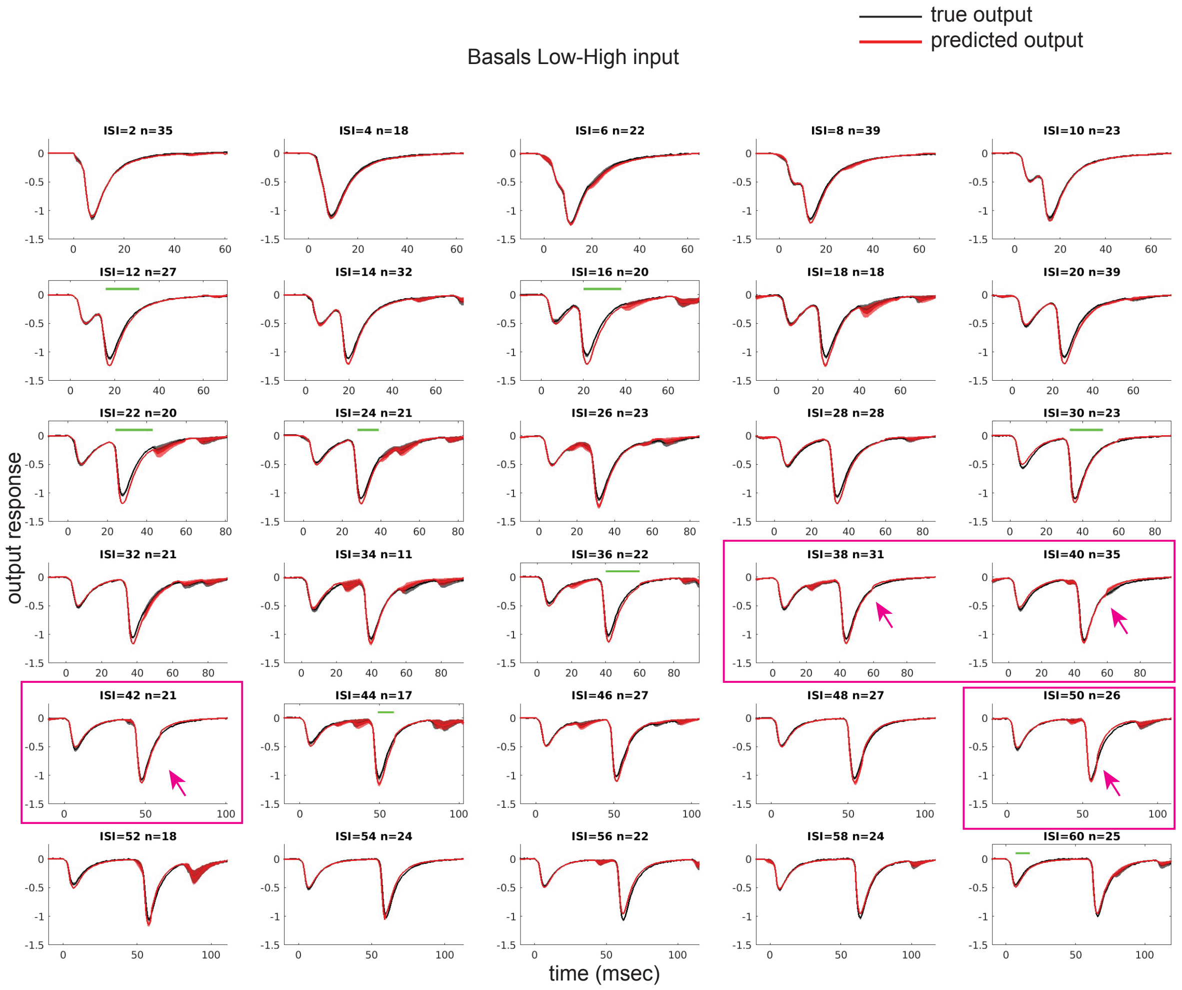


**Supplementary Figure 5. Paired pulse predictions for the basal dendrites (low magnitude followed by high magnitude paired pulses)**

Same as Supplementary Figure 4, but for low followed by high magnitude paired pulses. ISIs for which significant differences were found are indicated by black boxed plots (black square) and the time duration that significantly differed between the true and predicted traces is indicated with a green line.

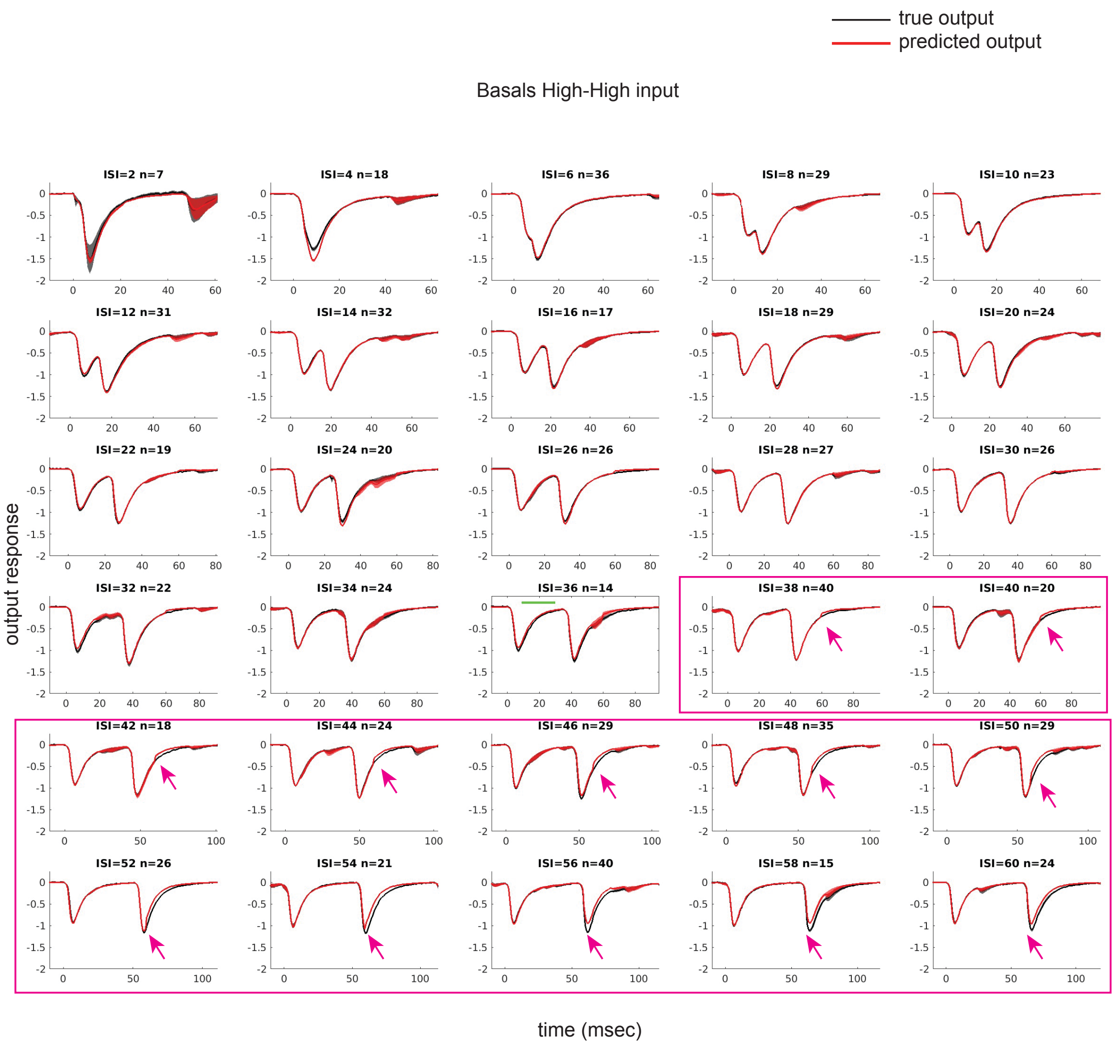


**Supplementary Figure 6. Paired pulse predictions for the basal dendrites (high magnitude for both pulses)**

Same as Supplementary Figure 4, but for high magnitude pulses.

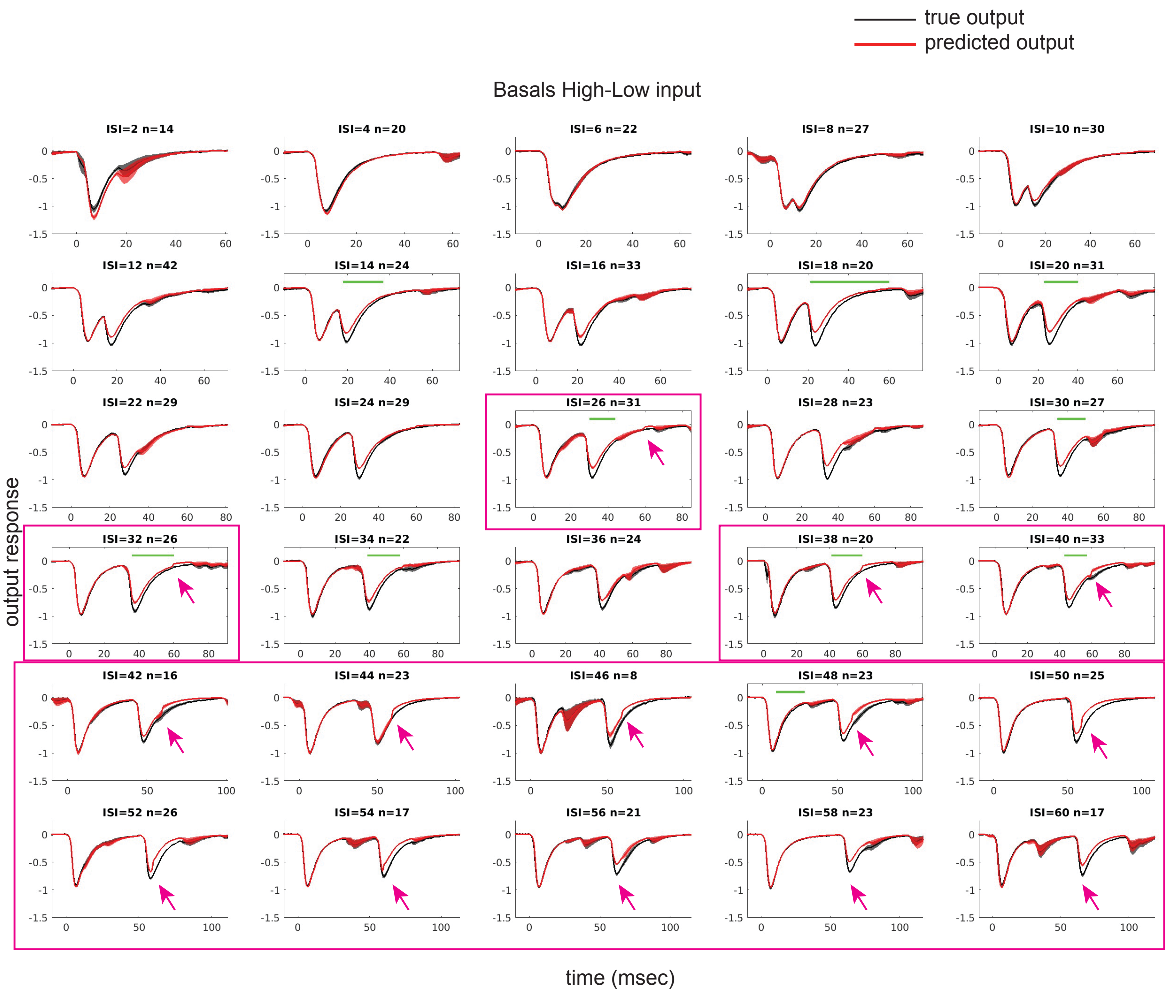


**Supplementary Figure 7. Paired pulse predictions for the basal dendrites (high magnitude followed by low magnitude paired pulses)**

Same as Supplementary Figure 4, but for high followed by low magnitude paired pulses.

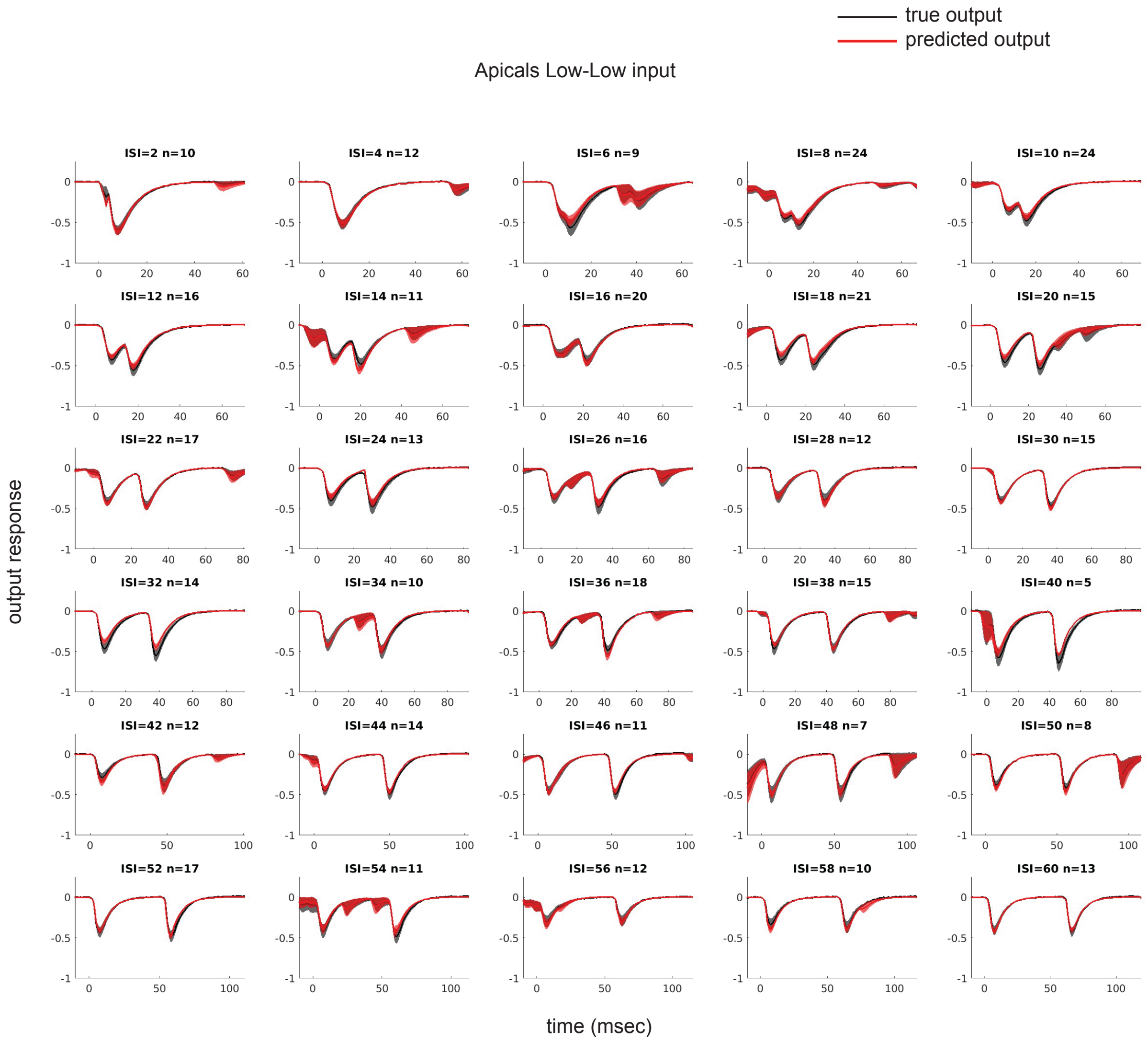


**Supplementary Figure 8. Paired pulse predictions for the apical dendrites (low magnitude for both pulses)**

Same as Supplementary Figure 4, but for the apical dendrites.

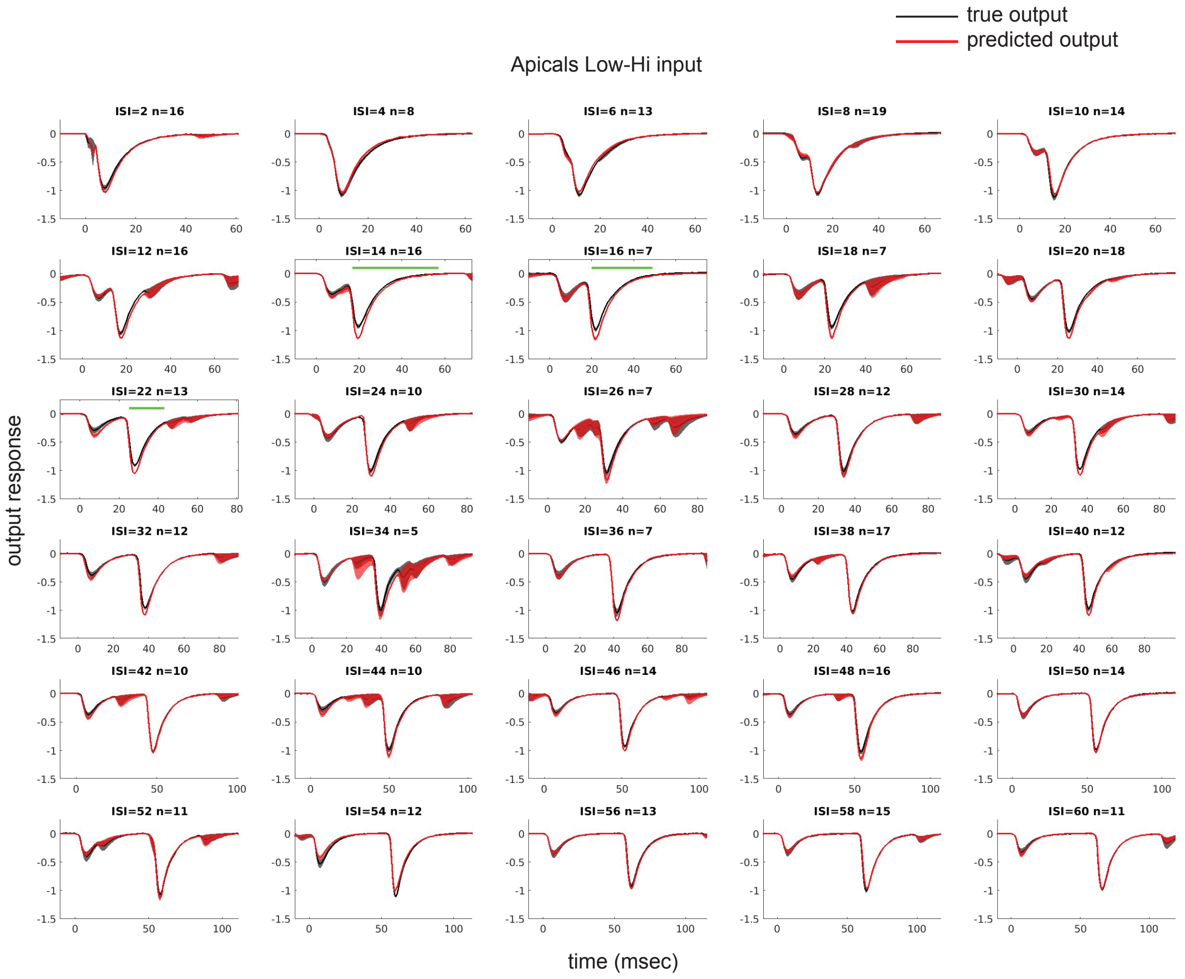


**Supplementary Figure 9. Paired pulse predictions for the apical dendrites (low magnitude followed by high magnitude paired pulses)**

Same as Supplementary Figure 5, but for the apical dendrites.

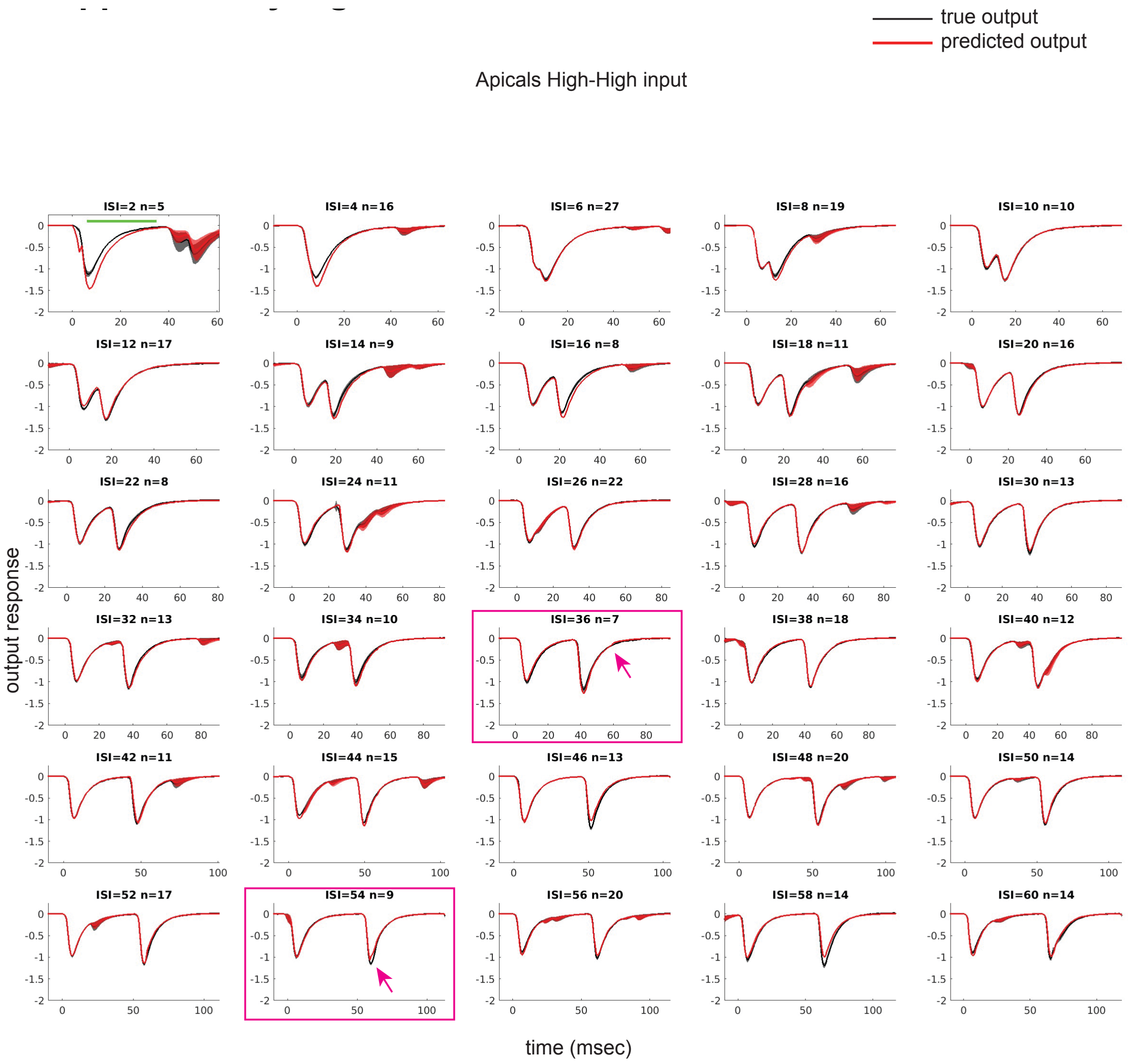


**Supplementary Figure 10. Paired pulse predictions for the apical dendrites (high magnitude for both pulses)**

Same as Supplementary Figure 6, but for the apical dendrites.

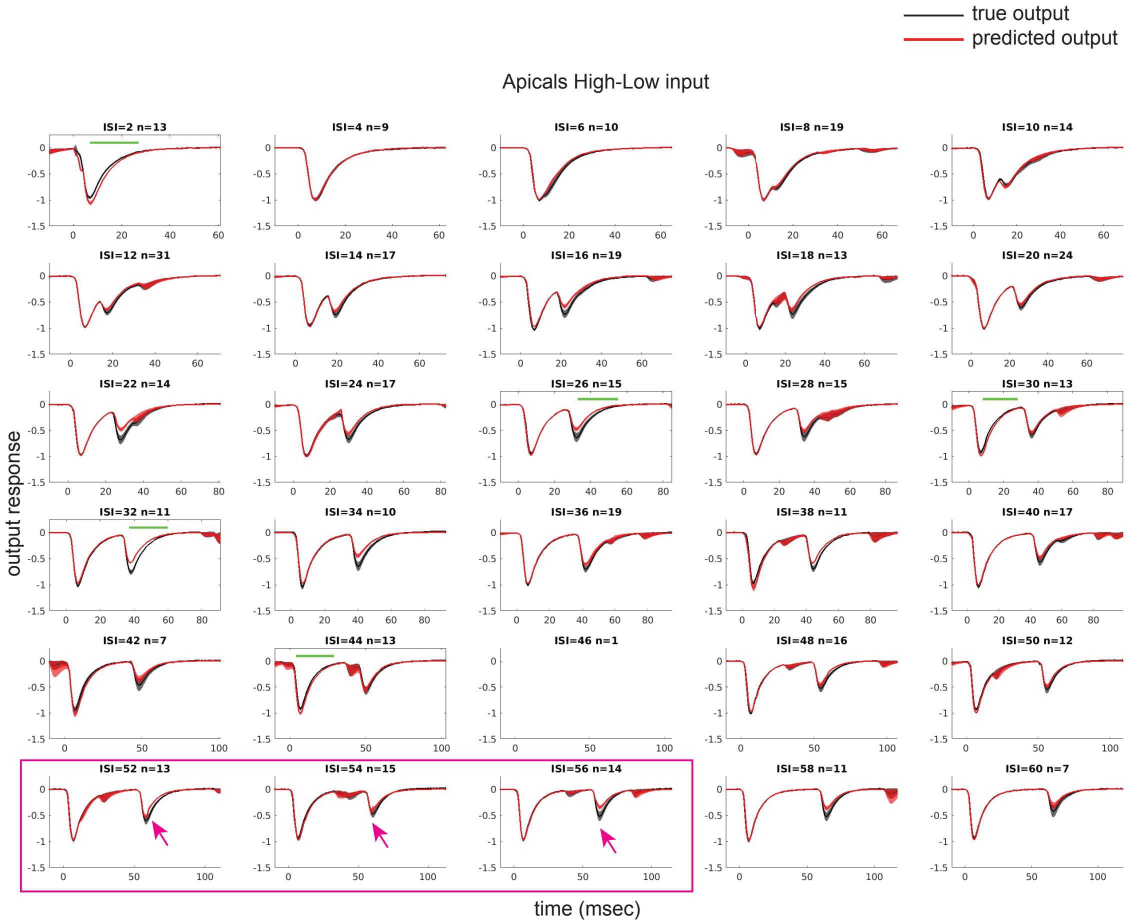


**Supplementary Figure 11. Paired pulse predictions for the apical dendrites (high magnitude followed by low magnitude paired pulses)**

Same as Supplementary Figure 7, but for the apical dendrites.

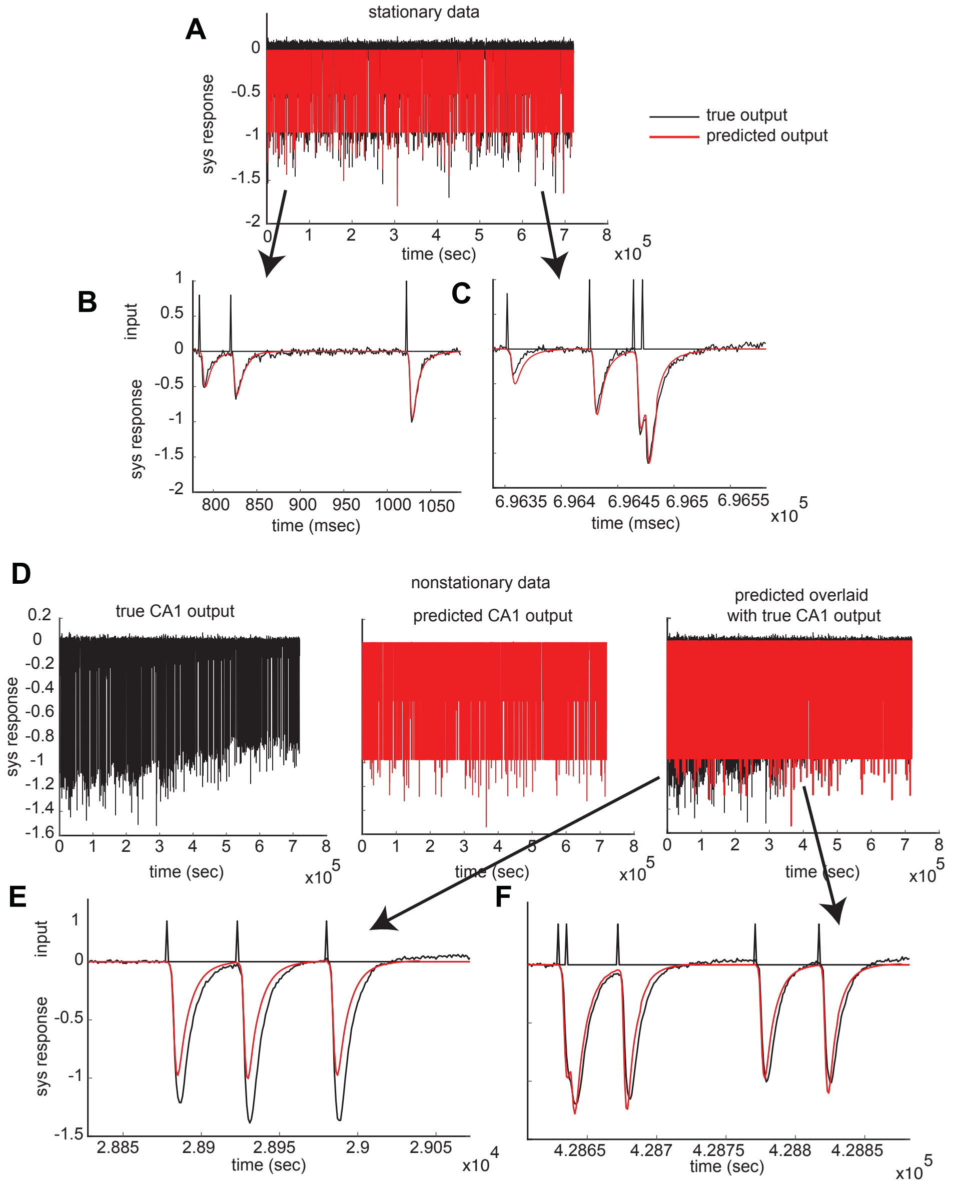


**Supplementary Figure 12. Nonstationarity is a source of prediction error**

**A,** Example of a stationary (constant response magnitude as a function of time) recording from the basal dendrites. Black and red traces reflect true and basal-transfer-function predicted responses, respectively. **B-C,** True output is within the range of the predicted output for segments extracted from both the beginning (B) and end (C) of the recording. **D,** Example of a nonstationary (decreasing response magnitude as a function of time) recording from the apical dendrites (left panel), predicted output (stationary by definition, middle panel), and both predicted and true outputs overlaid (right panel). Note the true outputs are outside (more negative) the prediction range, in the first half of the recording (right panel). **E-F,** True output is outside (E) and within (F) the range of the predicted output for segments extracted from the beginning (E) and middle (F) of the recordings, respectively. It is important to note that data was not excluded for nonstationary, for both model estimation and testing (see exclusion criteria list under ‘Analysis: kernel estimation’ section in the methods). The influence of nonstationarity was minimized analytically by indexing only the last 12 minutes of all 15-minute recording sessions for all data analyses.

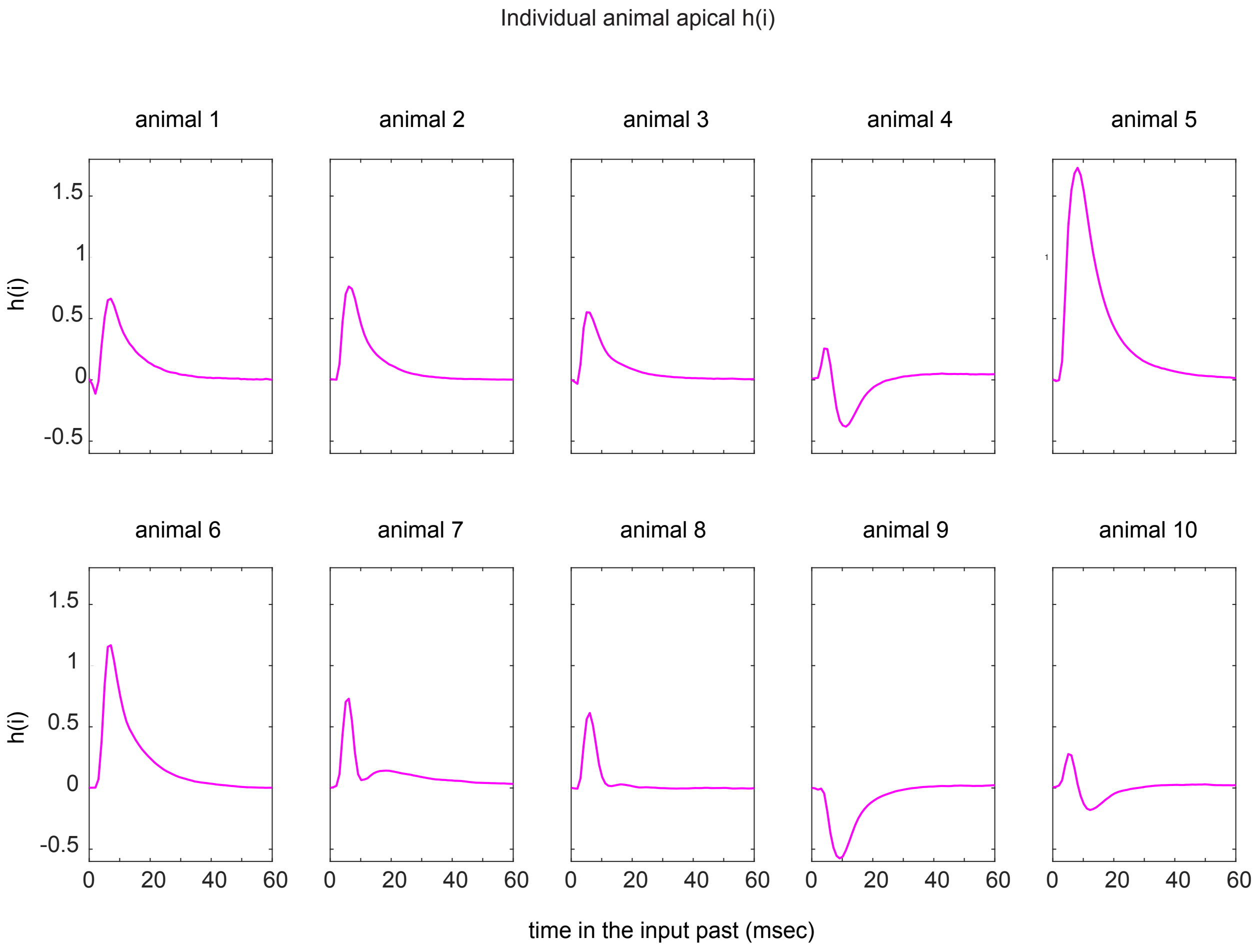


**Supplementary Figure 13. Individual animal *h(i)* estimates**

Each trace is an average across session (last 12-minutes) estimates recorded in the same animal (see Table 1).

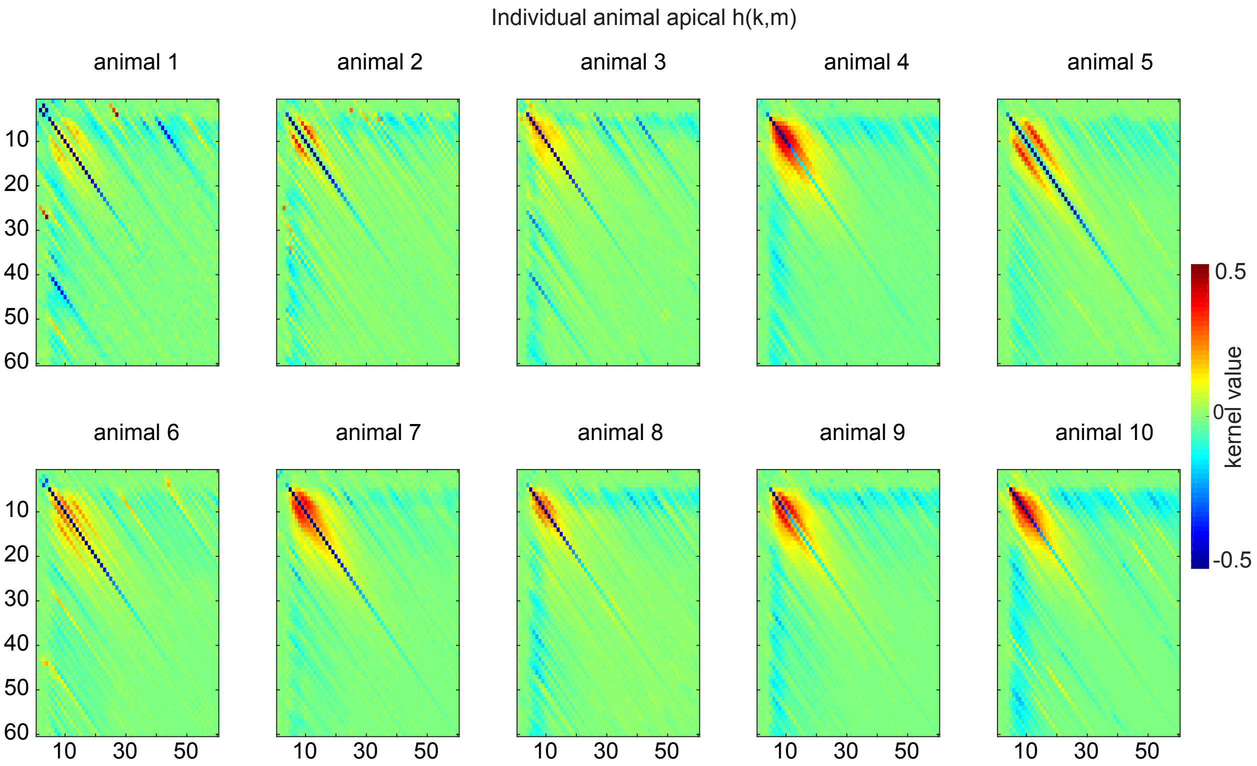


**Supplementary Figure 14. Individual animal *h(k,m)* estimates**

Each matrix is an average across session (last 12-minutes) estimates recorded in the same animal (see Table 1).

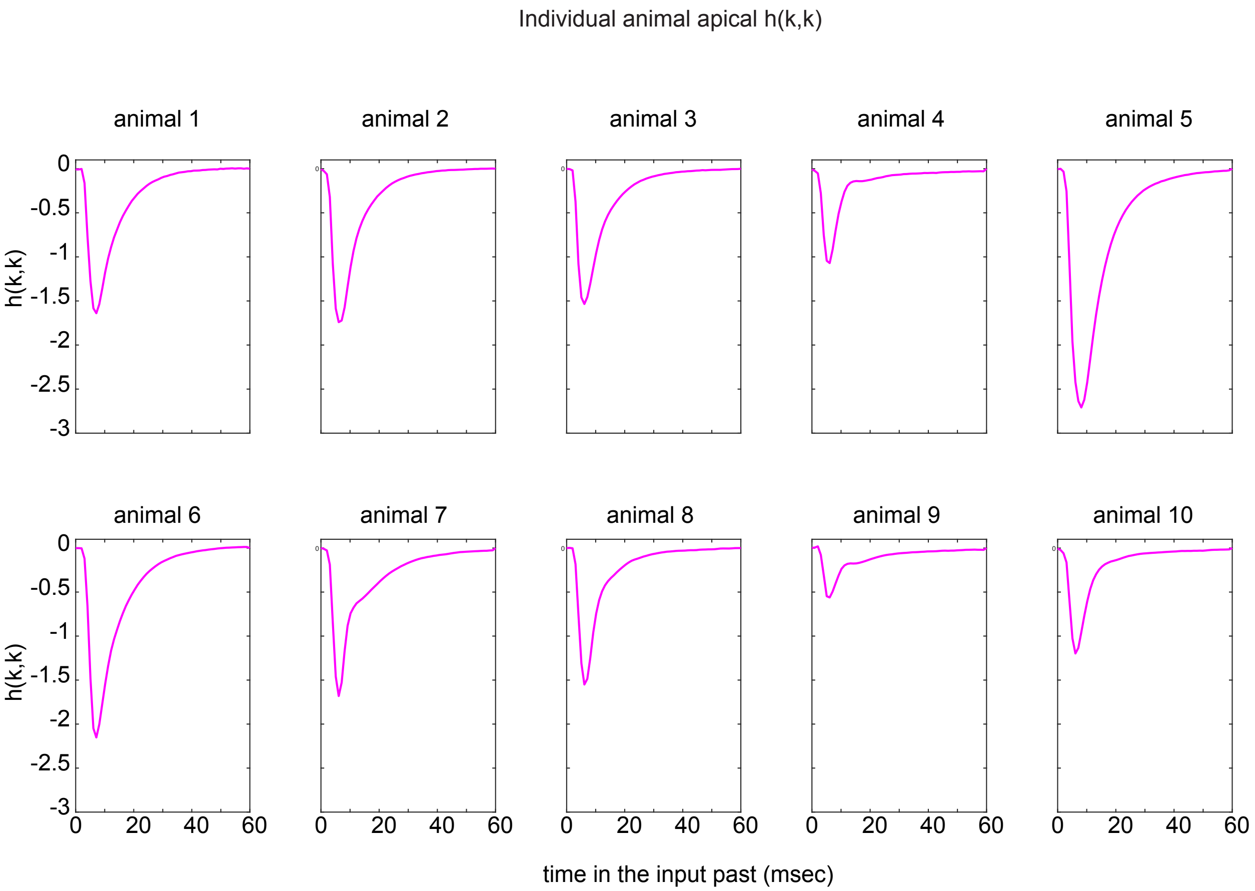


**Supplementary Figure 15. Individual animal *h(k,k)* estimates**

Each trace is an average across session (last 12-minutes) estimates recorded in the same animal (see Table 1).

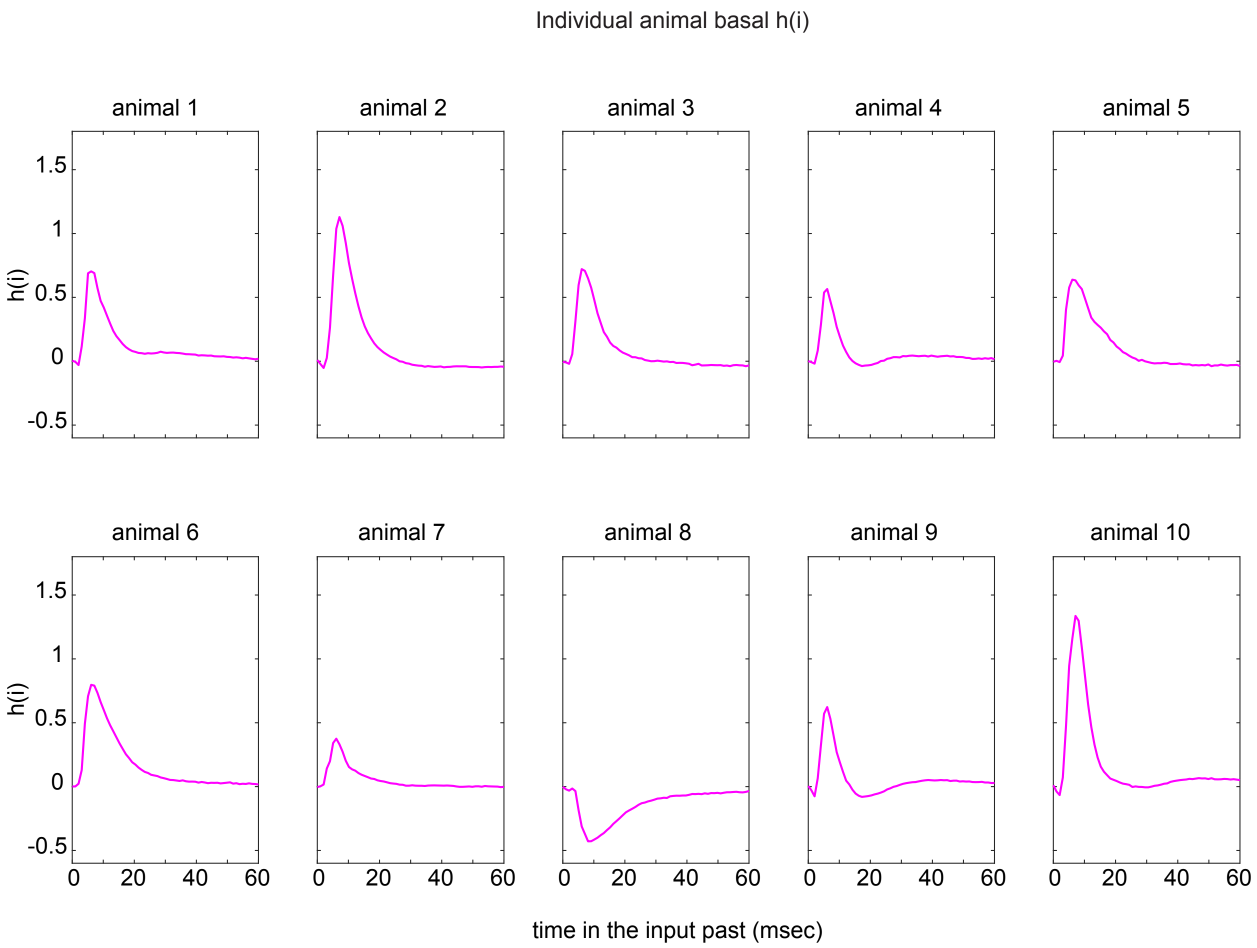


**Supplementary Figure 16. Individual animal *h(i)* estimates**

Each trace is an average across session (last 12-minutes) estimates recorded in the same animal (see Table 2).

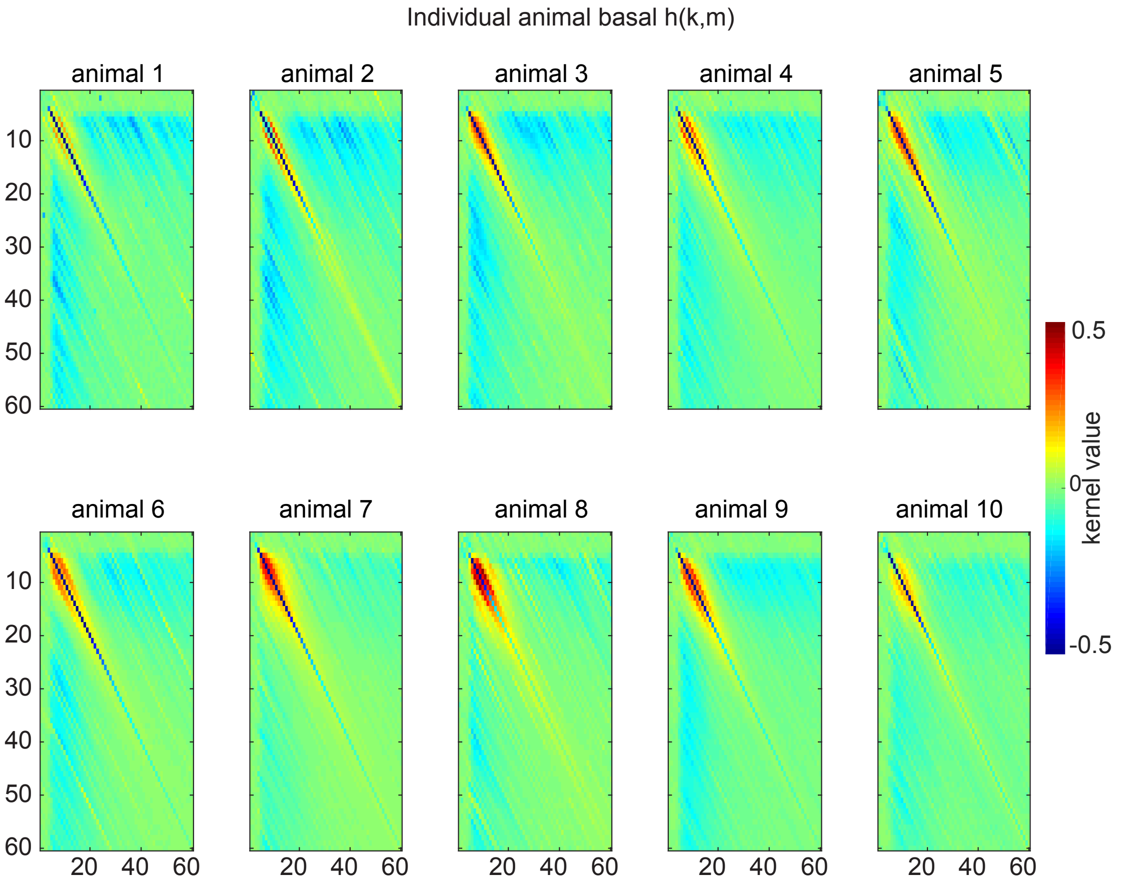


**Supplementary Figure 17. Individual animal *h(k,m)* estimates**

Each matrix is an average across session (last 12-minutes) estimates recorded in the same animal (see Table 2).

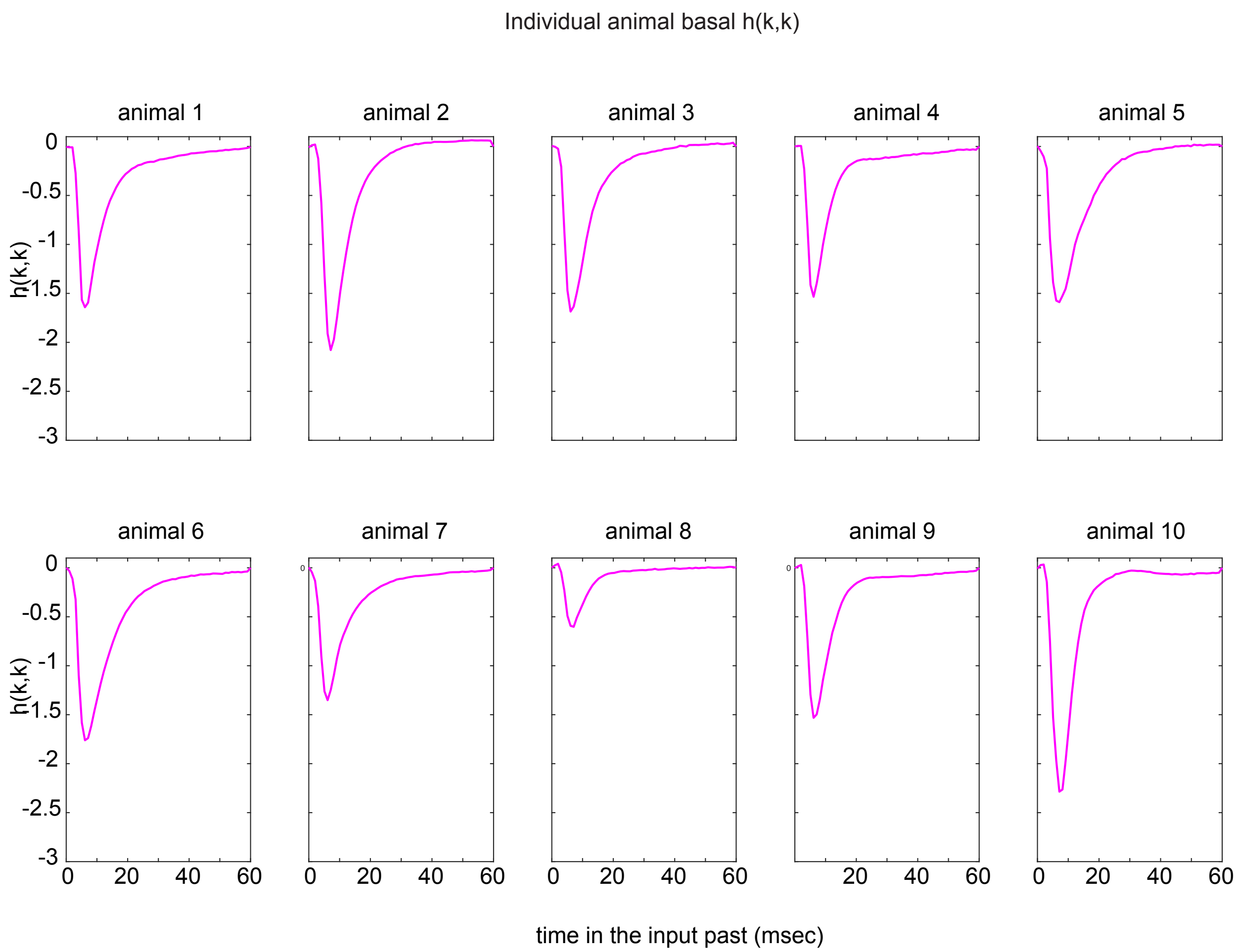


**Supplementary Figure 18. Individual animal *h(k,k)* estimates**

Each trace is an average across session (last 12-minutes) estimates recorded in the same animal (see Table 2).

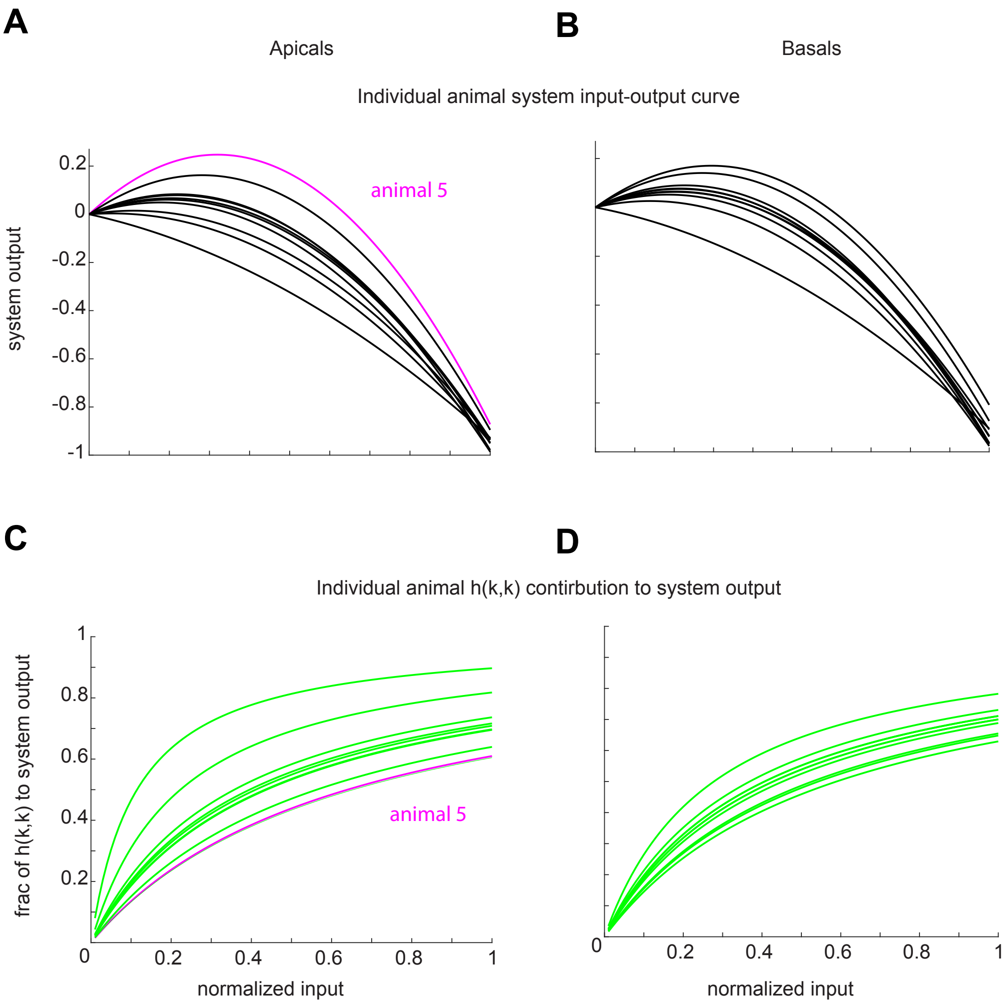


**Supplementary Figure 19. Individual animal input-output curves and nonlinear contribution to CA1 output**

**A-D,** Same as in Fig. 4A but plotted separately for (**A-B**) system output and (**C-D**) *h(k,k)* contribution to system output as a function of normalized input. Here, system output is defined as a univariate value (the peak value of the fEPSP) rather than the continuous fEPSP waveform. The timing of the peak ($7^{th}$ msec after stimulation) was identified using the group level *h(i)* peaks for apicals and basals (Fig. 4A and Fig. 6F, respectively). Left and right panels are for the apical and basal systems, respectively. Note the higher variability across animals for the apicals compared to the basals. Animal 5 is indicated in magenta as an example of different system input-output curves (top left panel) compared to the group. This is a consequence of a highly varied scaling of *h(i)* compared to other animal's estimates (see **Supplementary Figure 13**), and the fact that for this animal, system nonlinearity is only in effect for much larger normalized input values (rightward shift in bottom left panel); this means that such a varied *h(i)* will predominate the output for a larger range of input values, leading to a correspondingly varied input-output curve.
